# Geometric-Evolutionary Deep Learning Decodes the Human GPCR-Metabolome Interactome and Enables Systematic De-Orphanization

**DOI:** 10.64898/2026.02.21.707232

**Authors:** Tomoya Sakuma, Yuki Otani, Hideyuki Shimizu

## Abstract

G protein-coupled receptors (GPCRs) are the largest class of drug targets, yet hundreds of orphan GPCRs lack known endogenous ligands, limiting our understanding of human physiology and therapeutic development. Existing computational approaches often fail to generalize to these unseen targets due to a reliance on target-specific priors and linear feature integration. Here, we present G-LEAP, a deep learning framework that learns generalizable principles of GPCR-ligand recognition by synergizing evolutionary protein language models with 3D-aware geometric molecular representations. By implementing a bilinear interaction module, G-LEAP explicitly models non-linear cross-modal interactions and achieves superior generalization on stringent benchmarks, outperforming state-of-the-art methods by 22.4% in error reduction. Crucially, G-LEAP demonstrates robust chemical discrimination by effectively distinguishing active ligands from property-matched physicochemical decoys in the DUDE-Z benchmark and correctly rejecting 67% of hard-negative artifacts prioritized by physics-based docking simulations. Leveraging this capacity, we constructed a comprehensive atlas of over 120 million predicted interactions between 217,000 human metabolites and the GPCR superfamily, which ranked the true endogenous ligand within the top 1% of candidates for 33.7% of known pairs and identified putative orphan ligands validated by significant tissue-specific co-expression with their biosynthetic enzymes. Furthermore, large-scale virtual screening retrieved potent hits with novel chemical scaffolds distinct from known GPCR ligands, demonstrating robust scaffold hopping. G-LEAP thus provides a systematic and biologically validated platform to accelerate de-orphanization and expand the therapeutic chemical space.

## Introduction

G protein-coupled receptors (GPCRs) constitute the largest superfamily of transmembrane proteins, acting as the primary sensors that translate extracellular signals into intracellular responses. They represent the most successful class of therapeutic targets, accounting for over one-third of all FDA-approved medicines [1,2]. Their dysfunction is implicated in a vast array of human pathologies, ranging from neuropsychiatric disorders to cancer and metabolic diseases [3–5]. However, despite decades of extensive high-throughput screening and labor-intensive experimental de-orphanization efforts, over 120 GPCRs remain as “orphans” with undefined endogenous ligands and physiological functions, approximately 30% of the non-olfactory human GPCRome, leaving them effectively undrugged [6]. These orphan receptors constitute a vast “dark matter” in drug discovery, representing a critical knowledge gap that not only limits our understanding of human physiology but also severely hinders the development of next-generation therapeutics.

Computational approaches offer a promising alternative, yet existing methods face critical bottlenecks when applied as primary screening tools for GPCRs. While physics-based molecular docking remains valuable for post-hoc structural validation, it often fails for GPCRs due to the flexibility of their binding pockets and the lack of high-resolution structures for orphan targets [7,8]. Conversely, machine learning models have shown promise, but general protein-ligand interaction models, often biased toward soluble proteins, struggle to capture the unique properties of GPCRs and exhibit limited accuracy for this family [9]. To address this, specialized GPCR prediction methods have been developed [10–13], but these frequently rely on detailed structural annotations, such as specific ligand interaction sites within the binding pocket. Consequently, these models perform well on characterized families but fail to generalize to “unseen” orphan targets in the training data. Therefore, the critical challenge is to move beyond statistical pattern matching of sequences and instead develop a framework that comprehends generalizable patterns of molecular recognition transferable to unseen GPCR families.

To address this challenge, we developed G-LEAP (Graph-based Ligand Engine for Action Prediction), a deep learning framework specifically designed to decode GPCR-ligand recognition. Unlike conventional methods that rely on manual feature engineering, G-LEAP integrates evolutionary information from protein language models with 3D-aware geometric representations from molecular pretraining. These dual representations are synthesized through a bilinear interaction module, which models the cross-modal compatibility between the high-dimensional latent spaces. By explicitly modeling non-linear interactions, G-LEAP learns to discriminate active ligands based on structural complementarity rather than superficial physicochemical similarity, enabling it to distinguish true binders from property-matched decoys and reject “hard-negative” artifacts that frequently mislead physics-based simulations.

In this study, we demonstrate that G-LEAP achieves robust generalization to unseen receptor families, attaining an ROC-AUC of 0.829 ± 0.028 on the Protein-split benchmark and reducing error by 22.4% compared to state-of-the-art methods. Leveraging this capability, we applied G-LEAP to large-scale virtual screening and identified potent hits with novel chemical scaffolds exhibiting significant scaffold-hopping potential. Furthermore, we constructed a comprehensive atlas encompassing over 120 million predicted interactions between 217,000 human metabolites and the GPCR superfamily. The reliability of this atlas is validated by ∼34-fold enrichment in recovering known endogenous ligands and observing significant tissue-specific co-expression between predicted orphan receptor-ligand pairs and their biosynthetic enzymes. G-LEAP thus serves as a foundational resource to systematically accelerate the de-orphanization of the GPCR superfamily and expand the druggable chemical space without the prerequisite of target-specific structural priors.

## Results

### Decoding GPCR molecular recognition via geometric-evolutionary deep learning

To develop a robust framework for GPCR–ligand interaction prediction, we adopted a systematic and stepwise feature refinement strategy to quantify the specific contributions of evolutionary and geometric information to model performance. Utilizing datasets constructed from GLASS [14] and GPCRdb [15], we established a baseline using a conventional Graph Convolutional Network (GCN) [16] architecture trained with standard features such as protein mol2vec [17] and compound RDKit descriptors [18], based on the methodology of Zhang et al. [10]. This baseline achieved modest performance with ROC-AUC scores of 0.706 for the Random split and 0.701 for the Ligand split, while it showed notably lower performance of 0.687 on the challenging Protein split **(Supplementary Fig. 1A)**. The Protein split is particularly critical for simulating a true de-orphanization scenario because orphan GPCRs lack any prior ligand-binding data by definition and effectively represent “unseen” targets to the model. Consequently, achieving high predictive accuracy on this split serves as a direct proxy for the real-world capacity to identify the endogenous ligands for orphan GPCRs.

In the first refinement phase, we replaced the protein features with embeddings from the large-scale protein language model ESM Cambrian (ESM C) 300M (960 dimensions) [19]. This substitution yielded a remarkable performance leap across all metrics, where the Random split improved to 0.797 (+0.091), the Ligand split to 0.796 (+0.095), and the Protein split to 0.715 (+0.028) **(Supplementary Fig. 1A)**. Crucially, the substantial improvement in the Ligand split—where test compounds are distinct from training data—demonstrates that ESM C embeddings capture conserved binding site features that enable generalization to unseen compounds primarily through refined protein representations.

In the second phase, we replaced the compound features with an 881-dimensional representation combining embeddings from the pre-trained Uni-Mol2 model (768-dim) [20] and auxiliary RDKit descriptors (113-dim). This integration of 3D-aware geometric features resulted in further significant gains, leading to a Random split of 0.951 (+0.154) and a Ligand split of 0.937 (+0.141), while the Protein split surged to 0.822 (+0.107) **(Supplementary Fig. 1A)**. The dramatic improvement in the Protein split highlights the critical role of 3D molecular geometry, as Uni-Mol2 representations likely enable the model to infer binding feasibility based on steric and electrostatic complementarity. This capability is essential when predicting interactions for receptors not encountered during training. Together with the ESM C protein embeddings (960-dim) established in the first phase, this dual-modal feature set constitutes the optimal input representation for G-LEAP.

Having established high-quality feature representations, we next addressed the limitation of conventional GCN architectures that typically integrate protein and ligand features via simple concatenation [10]. This approach fails to explicitly model the complex and non-linear pairwise interactions necessary to capture cross-modal feature dependencies between the protein target and drug candidate. To overcome this, we introduced a bilinear interaction module to fuse the dual embeddings, which explicitly models the reciprocal compatibility between GPCRs and compounds rather than relying on linear fusion techniques. The integration of the bilinear interaction module into the GCN backbone resulted in consistent performance enhancements across all data splitting strategies compared to the concatenation baseline on the initial GLASS dataset, achieving ROC-AUC scores of 0.984 for Random split, 0.985 for the Ligand split, and 0.890 for the stringent Protein split. The consistent improvement in the Protein split is particularly significant as it demonstrates that explicit interaction modeling allows the network to capture generalizable binding patterns that transfer to novel receptors. Furthermore, among various Graph Neural Network architectures tested, the GCN-Bilinear configuration yielded the highest average performance across all splits and was thus established as the core architecture for G-LEAP **(Fig. 1A, Supplementary Table 1)**.

**Figure 1.**
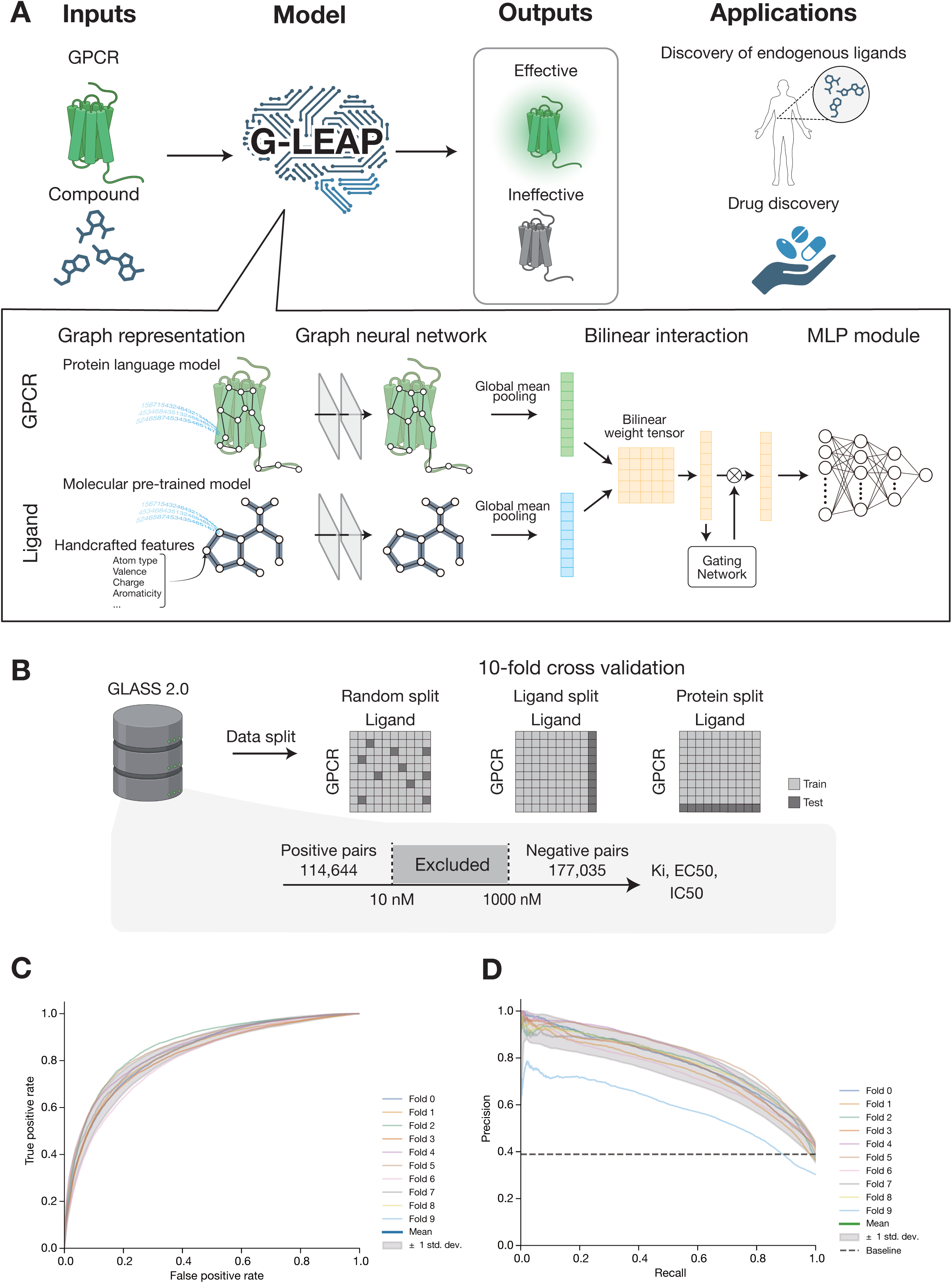
| The G-LEAP framework for GPCR-ligand interaction prediction. **A,** Conceptual overview and model architecture. G-LEAP integrates a GPCR sequence and a candidate compound (SMILES) to predict interaction probability by synergizing evolutionary protein context with 3D-aware molecular geometry. The receptor’s 3D structure predicted by AlphaFold is converted into a residue-level graph based on Cα–Cα spatial proximity (< 5 Å), with node features derived from ESM Cambrian embeddings. The ligand is represented as an atomic graph in which each node combines a 768-dimensional Uni-Mol2 embedding, capturing 3D conformational geometry, with 113-dimensional handcrafted physicochemical descriptors (atom type, valence, charge, aromaticity). Both 3D-informed graph representations are processed by independent Graph Neural Network (GNN) encoder streams and fused via a bilinear interaction module that explicitly models non-linear cross-modal interactions, followed by a multilayer perceptron (MLP) module for binary classification. **B,** Data processing and splitting strategy. The GLASS 2.0 dataset was rigorously filtered using gap-based activity thresholds (Positive: < 10 nM, Negative: > 1,000 nM) to exclude ambiguous data points. To strictly assess generalizability and prevent data leakage across receptors, 10-fold cross-validation was performed using a Protein split (where test proteins are unseen during training) in addition to standard Random and Ligand splits. The Protein split most stringently simulates the de-orphanization scenario, where test GPCRs are entirely absent from the training data. **C,** Receiver Operating Characteristic (ROC) curves for the Protein split 10-fold cross-validation. Individual fold curves (colored lines) and the mean ROC curve (blue line) with ±1 standard deviation (shaded area) demonstrate robust and consistent performance (mean AUC = 0.829 ± 0.028) across all folds, highlighting the model’s ability to generalize to novel, unseen receptors. **D**, Precision-Recall (PR) curves for the same Protein split cross-validation. The mean PR curve (green line) with ±1 standard deviation (shaded area) shows consistent predictive performance (mean Average Precision = 0.758 ± 0.062), confirming reliable identification of true positive GPCR–ligand interactions even under the stringent generalization setting.

#### Rigorous generalization capability and family-wise robustness

While the core GCN-Bilinear architecture was initially established through optimization on the original GLASS dataset, we sought to leverage the significantly expanded data volume of the GLASS 2.0 database, which was released in June 2025 [21]. This resource provided an approximately fourfold increase in data volume to 482,645 entries, enabling a more comprehensive training regime. We constructed a high-quality dataset comprising 291,679 pairs with 114,644 positive and 177,035 negative samples by applying strict activity thresholds of 10 nM and 1,000 nM respectively, excluding ambiguous intermediate values (**Fig. 1B**).

To maximize the potential of the GCN-Bilinear model on this expanded dataset, we conducted a systematic hyperparameter optimization across 97 combinations sweeping learning rates, weight decays, normalization strategies, and architectural parameters **(Supplementary Tables 2and 3)**. Sensitivity analyses revealed that G-LEAP maintains high performance across a broad range of embedding dimensions and learning rates **(Supplementary Fig. 2A)**, with batch sizes between 64 and 160 yielding comparable results **(Supplementary Fig. 2B)**. The optimized G-LEAP model achieved a statistically robust ROC-AUC of 0.829 ± 0.028 and Average Precision (AP) of 0.758 ± 0.062 on the rigorous Protein split **(Fig. 1C, D)**.

To benchmark G-LEAP against existing state-of-the-art methods, we retrained representative models including BIND [22], DEAttentionDTA [23], and Perceiver CPI [24] on the identical GLASS 2.0 dataset under the same experimental conditions. G-LEAP significantly outperformed all competitors, particularly on the Protein split where it achieved a ROC-AUC of 0.829 ± 0.028. This performance represents a 22.4% reduction in the error rate relative to the best-performing runner-up, Perceiver CPI (ROC-AUC of 0.780 ± 0.038) **(Table 1; Supplementary Tables 4and 5)**. The magnitude of this improvement on the Protein split, which is the most stringent generalization benchmark, provides compelling evidence that G-LEAP has acquired transferable representations of GPCR-ligand recognition applicable to structurally uncharacterized and orphan receptors.

**Table 1:** Performance comparison of G-LEAP and existing methods on the GLASS 2.0 dataset under the stringent Protein split.

| Method | ROC-AUC | Accuracy | Precision | Recall | F1 |
| --- | --- | --- | --- | --- | --- |
| BIND [22] | 0.634±0.063 | 0.629±0.120 | 0.543±0.233 | 0.287±0.258 | 0.312±0.193 |
| DEAttentionDTA [23] | 0.759±0.024 | 0.713±0.023 | 0.649±0.052 | 0.594±0.045 | 0.619±0.034 |
| Perceiver CPI [24] | 0.780±0.038 | 0.733±0.030 | 0.694±0.079 | 0.579±0.060 | 0.628±0.053 |
| **G-LEAP (Ours)** | **0.829±0.028** | **0.769±0.020** | **0.715±0.060** | **0.672±0.023** | **0.692±0.037** |

All methods were retrained and evaluated on the identical GLASS 2.0 dataset under the Protein split, where test receptors are entirely absent from the training data, simulating the de-orphanization scenario. Bold values indicate the best performance for each metric. Values show mean ± standard deviation over 10-fold cross-validation. Results for Random and Ligand splits are provided in Supplementary Tables 4 and 5, respectively.

We further evaluated the model’s robustness across the diverse GPCR superfamily using 10-fold cross-validation stratified by receptor class. Class A (Rhodopsin-like) GPCRs, which constitute 96.0% of the dataset and the majority of current drug targets, showed consistently high performance across all metrics **(Supplementary Fig. 3A–D)**. While performance variations were observed in data-limited classes such as Class B and Class C, this is an expected consequence of the severe data imbalance inherent to current GPCR databases. However, the model maintained discriminative power (ROC-AUC > 0.65) across all major families **(Supplementary Fig. 3B–D)**. Performance above this threshold confirms that G-LEAP has not overfit to the dominant Class A receptors but instead retains generalizable recognition capacity across the GPCRome. Together, these results establish G-LEAP as a robust and broadly applicable platform with particular utility for the therapeutically most relevant sectors of the GPCR superfamily.

#### Robust discrimination of active ligands from physicochemical decoys

A fundamental challenge in AI-driven drug discovery is enabling models to move beyond simple statistical correlations and learn to discriminate active ligands based on specific pharmacophoric features, rather than relying on superficial physicochemical proxies. To rigorously evaluate whether G-LEAP has achieved this capability, we challenged the model using the DUDE-Z benchmark [25]. This dataset is designed to be adversarial; for each active ligand, it contains multiple decoys that are meticulously matched for bulk physicochemical properties (e.g., molecular weight, LogP, charge) but lack biological activity (**Fig. 2A**). In this demanding test, G-LEAP demonstrated robust discriminative power, achieving strong enrichment of true ligands for three representative receptors. Specifically, for the D4 dopamine receptor (DRD4), the adenosine receptor A2a (ADORA2A), and the melatonin receptor type 1A (MTNR1A), the model reached ROC-AUC values of 0.963, 0.936, and 0.996 with corresponding AP scores of 0.594, 0.651, and 0.900, respectively **(Fig. 2B, C, Supplementary Fig. 4A)**.

**Figure 2.**
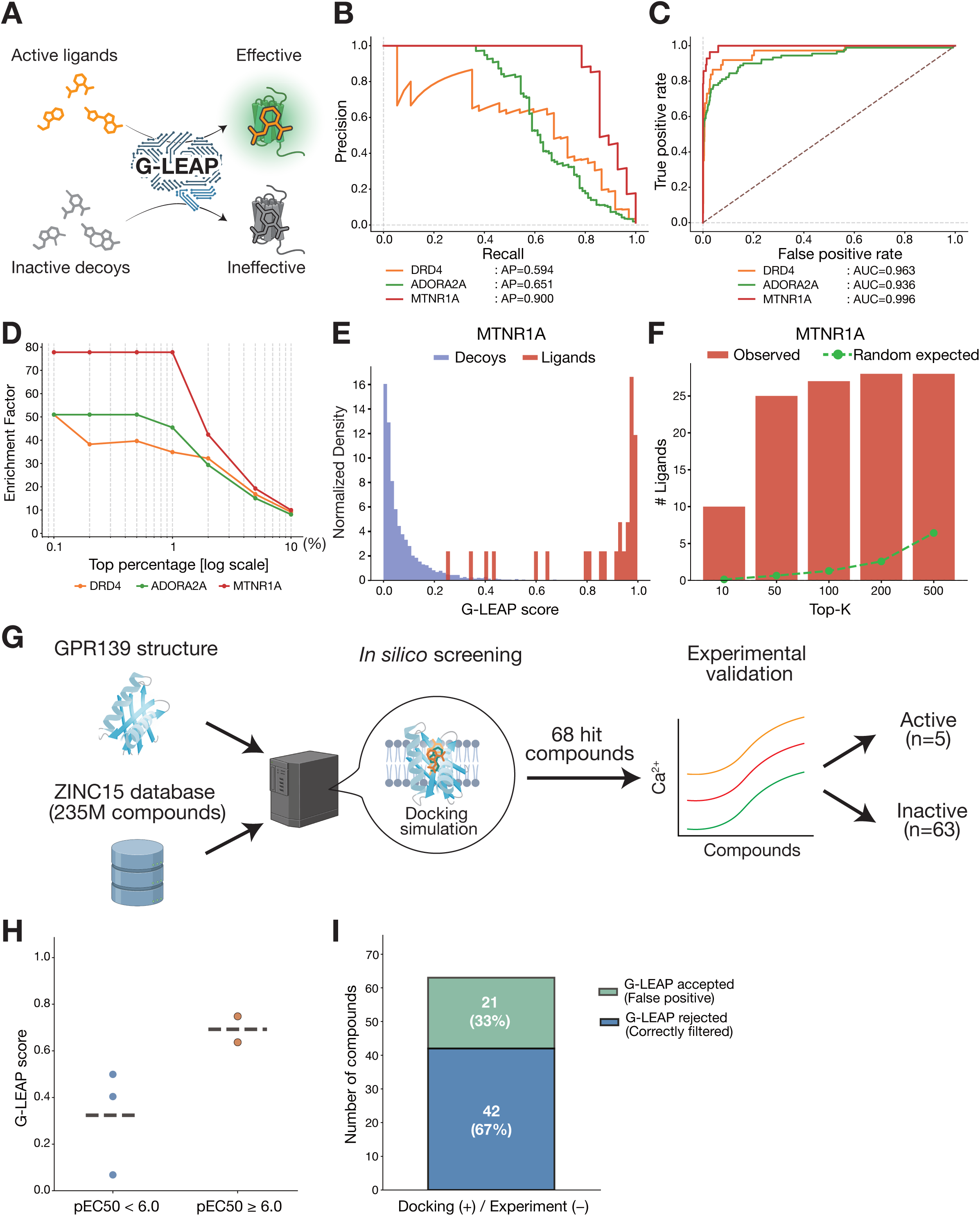
| Robust discrimination of active ligands from physicochemical decoys and external validation. **A**, Schematic overview of the discrimination task. G-LEAP was challenged to distinguish known active ligands from property-matched physicochemical decoys drawn from the DUDE-Z benchmark, in which decoys are carefully designed to match actives in bulk physicochemical properties (molecular weight, LogP, charge) while lacking biological activity. **B**, Precision-Recall (PR) curves for three GPCR targets from the DUDE-Z dataset: D4 dopamine receptor (DRD4), adenosine receptor A2a (ADORA2A), and melatonin receptor type 1A (MTNR1A). The legend reports the average precision (AP) for each target. **C**, ROC curves for the same three GPCR targets (DRD4: ROC-AUC = 0.963; ADORA2A: ROC-AUC = 0.936; MTNR1A: ROC-AUC = 0.996). The dashed diagonal line indicates the performance expected under random classification. **D**, Enrichment factor (EF, y-axis) curves as a function of the top-screened percentage (log scale, x-axis) for DRD4, ADORA2A, and MTNR1A. EF at a given top-K% is defined as the ratio of the observed number of actives retrieved to the number expected under random selection. All three targets show substantial early enrichment well above random expectation. **E**, The distribution of G-LEAP prediction scores for MTNR1A, comparing known active ligands (red) and decoys (blue). Histograms are normalized to unit area to enable comparison despite the large sample size disparities between groups (ligands: n = 28; decoys: n = 2,150). **F**, Numbers of active ligands retrieved at top-K% for MTNR1A. Observed counts (red bars) substantially exceed the number expected under random ranking (green dashed line) at all examined thresholds, confirming the practical screening utility of G-LEAP for this target. **G,** External validation workflow using an independent GPR139 agonist discovery dataset from Cabeza de Vaca *et al.* [28]. This independent test set contains hit compounds originally prioritized by physics-based docking (DOCK3.7). See Methods for details. **H,** Discrimination of compound potency by G-LEAP. Predicted scores are shown for GPR139 compounds tested in Ca²⁺ mobilization assays, stratified by potency class: higher-potency (pEC₅₀ ≥ 6.0) received higher G-LEAP scores than lower-potency compounds (pEC₅₀ < 6.0), demonstrating a positive correlation between predicted score and experimentally measured activity. **I,** Rejection of false-positive docking hits. Of 63 compounds prioritized by DOCK3.7 (physics-based docking) but confirmed experimentally inactive (hard-negatives), G-LEAP correctly assigned low prediction scores to 42 (67%), highlighting its capacity to filter structure-based false positives that arise from approximations in solvation energy and entropic penalty.

We next computed enrichment factors (EF) at top-K% thresholds and found that G-LEAP exhibited substantial early enrichment well above random expectation (**Fig. 2D, Supplementary Fig. 4B**). Notably, the G-LEAP scores were clearly separated between ligands and decoys (**Fig. 2E**), with MTNR1A reaching an EF@1% of approximately 78 and maintaining high enrichment across broader top-K% ranges (**Fig. 2F**). DRD4 and ADORA2A also showed strong enrichment (EF@1% ∼51 and ∼40, respectively), indicating screening performance consistent with practical drug discovery needs [26,27] (**Supplementary Fig. 4A, B**). This consistent performance across multiple receptors suggests that G-LEAP is sensitive to the precise three-dimensional arrangement of pharmacophoric features required for receptor activation, rather than simply recapitulating global physicochemical properties.

We further validated the model’s practical utility using an independent, experimentally derived dataset from a recent GPR139 agonist discovery campaign **(Fig. 2G)** [28]. Unlike curated benchmarks, this dataset reflects the noisy reality of early-stage drug discovery: of 68 compounds originally prioritized by physics-based docking (DOCK3.7) [29], only five showed potent agonist activity in Ca²⁺ mobilization assays (EC₅₀: 160 nM–3.6 µM, pEC₅₀ 5.4–6.8), while the remaining 63 failed to yield validated activity. G-LEAP assigned higher predicted probabilities to compounds with higher potency (pEC₅₀ ≥ 6.0) than to those with lower potency (pEC₅₀ < 6.0) (**Fig. 2H**). Most notably, we assessed the model’s ability to reject false-positive docking hits: of the 63 compounds that had been highly ranked by DOCK3.7 but proved experimentally inactive, G-LEAP correctly assigned low prediction scores to 42 (67%) (**Fig. 2I**). Structure-based docking methods are known to produce false positives due to approximations in solvation energy and entropic penalty estimation [30]. The ability of G-LEAP to reject the majority of these compounds suggests that training on large-scale evolutionary and chemical data enables the model to implicitly account for biophysical factors that mislead physics-based simulations, thereby serving as an effective orthogonal filter to purify virtual screening hits.

#### Validation of screening utility and scaffold-hopping capacity on ligand-characterized GPCRs

Prior to applying G-LEAP to orphan GPCRs, we evaluated its screening utility through a simulated drug-discovery campaign against GPCRs with well-characterized ligands (**Supplementary Fig. 5A**). For this large-scale screening, we used 6,151,237 drug-like compounds retrieved and filtered from the ZINC20 database [31] (see Methods). As targets, we selected three representative GPCRs including the free fatty acid receptor 1 (FFAR1), the muscarinic acetylcholine receptor M2 (CHRM2), and the orexin receptor type 2 (HCRTR2).

To verify that G-LEAP does not assign spuriously high scores indiscriminately, we first analyzed the distribution of G-LEAP scores for each target (**Supplementary Fig. 5B**). The resulting distributions were non-uniform, with scores remaining low for the vast majority of compounds, indicating that G-LEAP infers binding feasibility for only a small, target-specific subset of the chemical library. To further characterize the chemical landscape of the retrieved compounds, we partitioned them into four groups comprising known ligands, high-scoring compounds, low-scoring compounds, and randomly selected compounds. UMAP analysis revealed that high-scoring compounds occupied a chemical space more closely resembling that of known ligands compared to low-scoring and randomly selected compounds (**Supplementary Fig. 5C**), supporting the ability of G-LEAP to preferentially retrieve bioactive compounds.

A key limitation of conventional ligand-based virtual screening is its reliance on structural similarity to known active compounds, which restricts exploration to chemically adjacent regions of drug space and can perpetuate intellectual property constraints [32,33]. To assess whether G-LEAP can overcome this limitation, we quantified the structural novelty of its top-ranked candidates relative to known ligands. We first established fingerprint-specific similarity thresholds based on pairwise Tanimoto distributions of random compound pairs (**Supplementary Fig. 6A**; Morgan cutoff: 0.25, RDKit cutoff: 0.50, MACCS cutoff: 0.65), and then measured the maximum Tanimoto similarity of each top-ranked candidate to any known ligand for each target (**Supplementary Fig. 6B**). Across all three GPCR targets, a substantial proportion of high-scoring compounds fell below these similarity thresholds, demonstrating that G-LEAP successfully retrieves structurally novel scaffolds markedly distinct from all known ligands. This scaffold-hopping capacity positions G-LEAP as a versatile framework for accessing underexplored regions of chemical space that would be unreachable through conventional similarity-based search approaches, with implications for expanding the accessible regions of therapeutic chemical space.

### Large-scale virtual screening identifies candidate ligands for clinically relevant orphan GPCRs

To demonstrate the practical utility of G-LEAP in a real-world drug discovery context, we conducted a large-scale virtual screening campaign targeting three clinically significant orphan GPCRs: GPR3, GPR17, and GPR88, which lack approved therapeutics despite associations with neurological disorders, inflammatory conditions, and brain and behavioral function, respectively [34–36]. We utilized the same ZINC20-derived library of 6,151,237 drug-like compounds described previously, filtered through a medicinal chemistry cascade comprising Lipinski’s rule of five [37], PAINS exclusion [38], and Brenk’s structural alerts [39] (**Fig. 3A**).

**Figure 3.**
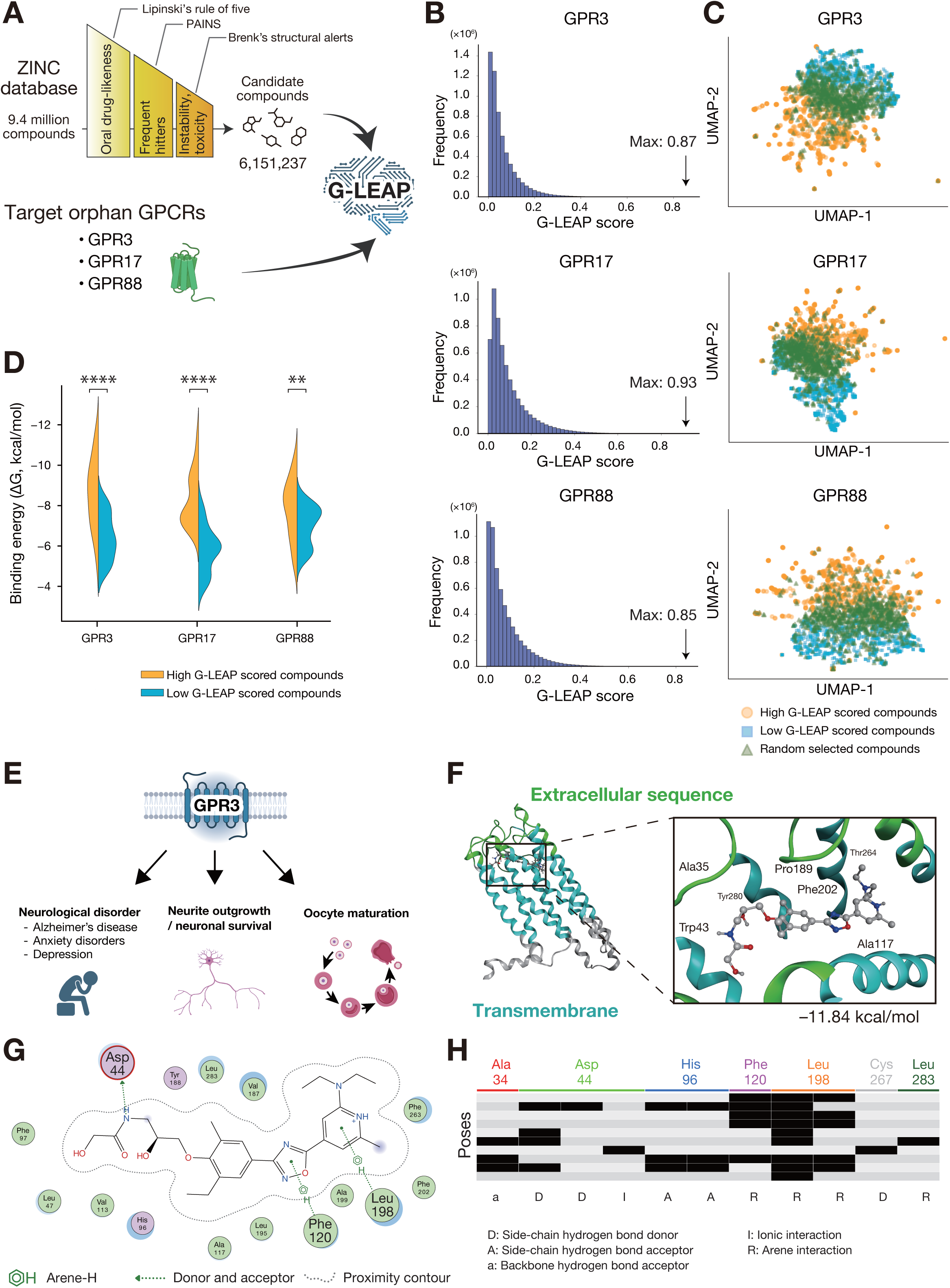
| Virtual screening of orphan GPCRs yields structurally diverse candidates with predicted binding modes. **A**, Large-scale virtual screening pipeline. The ZINC20 database (9.4 million compounds) was subjected to a medicinal chemistry filtration cascade comprising Lipinski’s rule of five [37], PAINS exclusion [38], and Brenk’s structural alerts [39], resulting in 6.15 million high-quality drug-like candidates. These compounds were screened against three clinically relevant orphan GPCRs that currently lack approved drugs targeting them: GPR3 (implicated in neurological disorders), GPR17 (inflammatory signaling), and GPR88 (brain and behavioral function) [34–36]. **B,** Distribution of G-LEAP prediction scores for GPR3, GPR17, and GPR88. Histograms display the frequency of scores across the 6.15 million screened compounds for each orphan receptor. The distinct, receptor-specific score distributions—with the majority of compounds receiving low scores—demonstrate that G-LEAP generates target-dependent predictions rather than indiscriminately assigning high scores. **C,** Chemical space analysis via UMAP projection. For each orphan GPCR target, compounds were partitioned into three groups: top-ranked 1,000 compounds (orange circles), bottom-ranked 1,000 compounds (blue squares), and 1,000 randomly selected compounds (green triangles). **D,** Predicted binding affinity comparison. Violin plots compare the predicted binding free energies (ΔG, kcal/mol) estimated by Boltz-2 for the top-ranked G-LEAP hits (High G-LEAP scored compounds, orange) versus low-scoring randomly selected compounds (Low G-LEAP scored compounds, blue). *P*-values were calculated using the two-sided Mann-Whitney *U* test (**: *p* < 0.01, ****: *p* < 0.0001). Top-ranked compounds consistently exhibited more favorable (more negative) predicted binding free energies across all three orphan targets, indicating strong concordance between G-LEAP’s predictions and physics-based energy landscapes. **E,** Biological context of GPR3. Schematic summary of reported physiological and pathological associations of GPR3, including roles in neurological disorders [41,42], neurite outgrowth and neuronal survival [43], and oocyte maturation [44,45], highlighting the therapeutic relevance of this orphan GPCR. **F,** Representative predicted binding mode of a top-scoring compound for GPR3. The Boltz-2–predicted structure shows the ligand positioned within the receptor binding pocket, with key residues in the transmembrane helices and extracellular loops highlighted. Predicted binding free energy: −11.84 kcal/mol. **G,** Two-dimensional ligand–receptor interaction diagram for the representative pose shown in panel F. Residues engaged in proximity contacts are enclosed in contour lines. **H,** Protein–ligand interaction fingerprint (PLIF) analysis across multiple predicted binding poses. Each row represents an independent Boltz-2 prediction; each column represents a binding-pocket residue. Black cells indicate the presence of a specific ligand–residue interaction (hydrogen bond, ionic, or arene contact); white cells indicate absence of interaction. The consistent reproduction of arene–H interactions across independent calculations (e.g., residues Phe120, Leu198) demonstrates that these contacts represent robust structural features of the predicted binding mode.

To confirm that G-LEAP was generating target-specific predictions rather than indiscriminately assigning high scores, we first examined the distribution of prediction scores for each orphan GPCR. The histograms revealed distinct, receptor-specific score profiles (**Fig. 3B**), with the vast majority of compounds receiving low scores and only a small subset achieving high ranks. This observation demonstrates that G-LEAP infers binding feasibility in a target-dependent manner. To characterize the chemical diversity of retrieved candidates, we performed UMAP projection of top-ranked compounds, low-ranked compounds, and randomly selected compounds. High-scoring candidates formed distinct clusters in chemical space, clearly separated from the random background of the ZINC database (**Fig. 3C**). The chemical space occupied by these candidates resembled the distributions observed for ligand-characterized GPCRs (**Supplementary Fig. 5C**), suggesting that G-LEAP retrieves compounds with pharmacologically relevant structural features even in the absence of known reference ligands.

To assess whether the high-scoring predictions correspond to physically plausible binding modes, we reconstructed 3D receptor–ligand complexes for top-ranked candidates using Boltz-2 [40], a state-of-the-art model for structure prediction and binding affinity estimation. Comparative analysis of top-ranked G-LEAP hits versus low-scoring randomly selected compounds revealed that the former group consistently exhibited more favorable predicted binding free energies (**Fig. 3D**), indicating strong concordance between the G-LEAP score and physics-based energy landscapes. For GPR3, an orphan GPCR implicated in several conditions, such as neurological disorders [41,42], neurite outgrowth and neuronal survival [43], and oocyte maturation [44,45], the predicted binding poses of top hits revealed extensive molecular recognition features including hydrogen bonds and hydrophobic contacts with key transmembrane residues (**Fig. 3E, F**). A two-dimensional interaction diagram illustrated the spatial arrangement of these contacts, highlighting arene interactions mediated by the ligand’s aromatic scaffold (**Fig. 3G**). To assess the robustness of the predicted binding mode, we performed protein–ligand interaction fingerprint (PLIF) analysis [46] across multiple docking poses and found that specific interactions, such as arene–H interactions with key residues including Phe120 and Leu198, were consistently reproduced across independent calculations (**Fig. 3G, H**). Taken together, these findings provide convergent computational evidence that the top-ranked G-LEAP predictions represent structurally realistic receptor–ligand complexes, offering a prioritized set of candidates for experimental validation and accelerating the discovery pipeline for orphan GPCR pharmacology.

#### A biologically validated interaction atlas of the human metabolome and GPCRs

One of the most significant barriers in GPCR pharmacology is the existence of hundreds of orphan receptors whose endogenous ligands remain unknown. Leveraging the robust generalization capability of G-LEAP to unseen targets, we sought to bridge this knowledge gap by constructing a comprehensive GPCR-Metabolome Atlas. We screened the entire Human Metabolome Database (HMDB) [47], comprising approximately 217,000 endogenous and exogenous metabolites, against the human GPCR superfamily. This large-scale computational campaign generated a global catalog of over 120 million putative ligand–receptor pairs (**Fig. 4A**).

**Figure 4.**
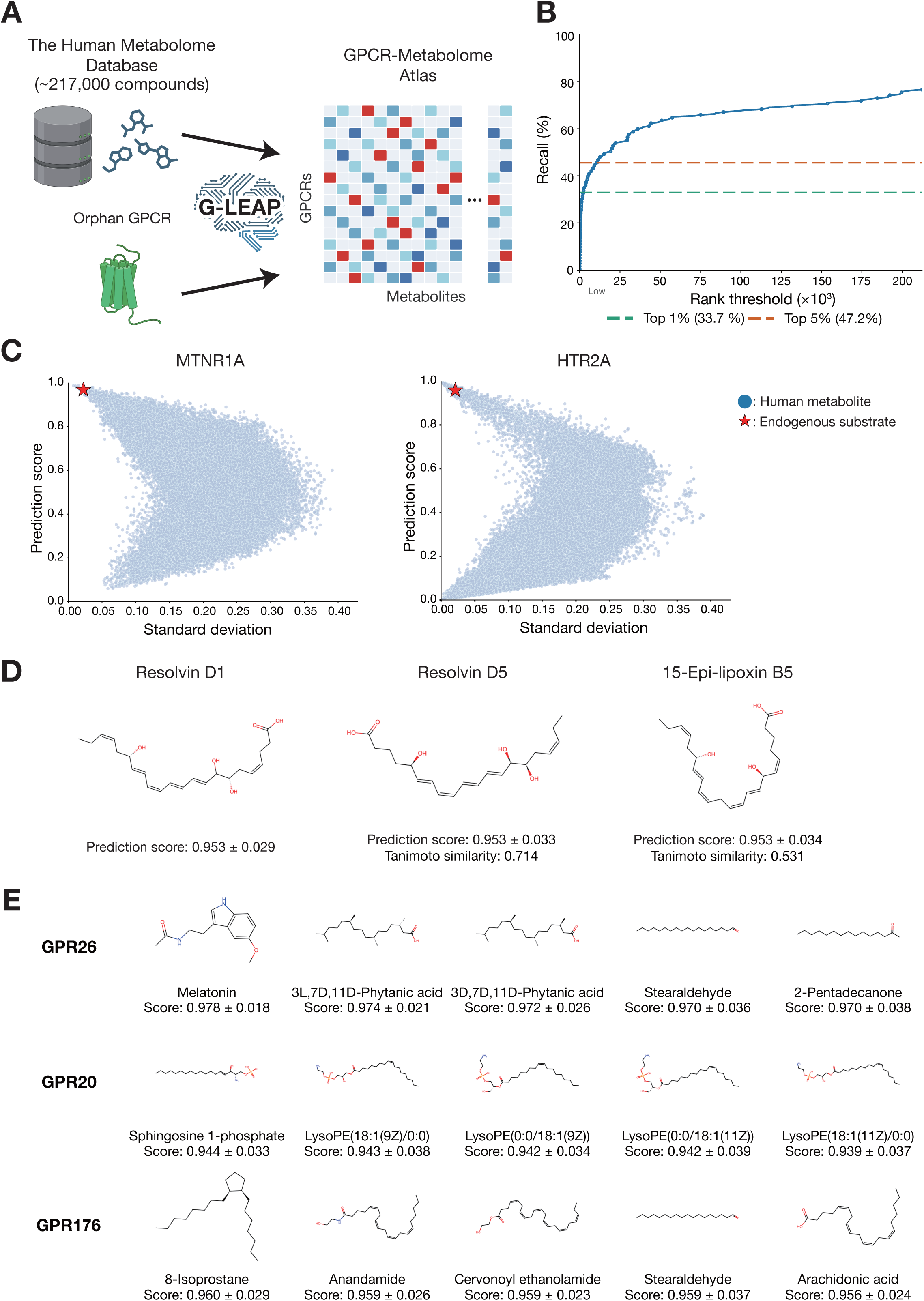
| A biologically validated interaction atlas of the human metabolome and GPCRs. **A**, Atlas construction workflow. The Human Metabolome Database (HMDB; ∼217,000 compounds) was screened against the human GPCR superfamily using G-LEAP, generating over 120 million putative endogenous ligand–receptor interaction predictions to construct a global atlas. **B,** Recall performance on known endogenous ligand–receptor pairs. Using 252 annotated pairs from the IUPHAR/BPS Guide to Pharmacology as ground truth, G-LEAP successfully retrieves the correct ligand within the top 1% of the ranked metabolome list for 33.7% of test cases (known pairs), and within the top 5% for 47.2% of cases. This represents a ∼34-fold enrichment over random selection. **C,** Validation with well-characterized receptor–ligand pairs. Scatter plots show G-LEAP prediction scores (y-axis) versus cross-validation standard deviation (CV-SD, x-axis) for all ∼217,000 HMDB metabolites screened against melatonin receptor type 1A (MTNR1A, left), and serotonin receptor 2A (HTR2A, right). Blue dots represent individual metabolites. The known endogenous ligands (melatonin and serotonin) are indicated as red stars positioned at the top-left of each plot, exhibiting high prediction scores (> 0.95) and low uncertainty (CV-SD < 0.05). This demonstrates G-LEAP’s ability to correctly prioritize true ligands from large metabolite libraries. **D,** Structural specificity of GPR32, for which Resolvin D1 has been proposed as a candidate endogenous ligand [49]. Chemical structures are shown for Resolvin D1 (left; prediction score: 0.953 ± 0.029) and structurally related lipid mediators enriched among top-ranked candidates: Resolvin D5 (middle; score: 0.953 ± 0.033; Tanimoto similarity to Resolvin D1: 0.714) and 15-Epi-lipoxin B5 (right; score: 0.953 ± 0.034; Tanimoto similarity: 0.531). Tanimoto similarity scores reflect chemical relatedness to Resolvin D1, confirming G-LEAP extracts pharmacophore-like features aligned with structural constraints of lipid-binding GPCRs. **E,** High-confidence predictions for orphan receptors. Top-ranked metabolite candidates with high prediction scores and low uncertainty for three representative Class A orphan receptors, namely GPR26, GPR20, and GPR176. These predictions align with phylogenetic relationships and provide testable hypotheses for experimental de-orphanization efforts.

To ensure the physiological relevance of this atlas, we refined our prediction strategy specifically for endogenous interactions. We redefined the activity threshold to 10 µM, which reflects the typical affinity range of endogenous ligands and is substantially weaker than the nanomolar affinities commonly observed for synthetic drugs [48]. Furthermore, we employed an ensemble scoring approach derived from the 10-fold cross-validation models. This method not only improved robustness on new data (**Supplementary Fig. 7A**) but also enabled us to quantify prediction reliability using the cross-validation standard deviation (CV-SD) as an uncertainty metric.

We quantified the systematic reliability of the atlas using a rigorous recall analysis based on the IUPHAR/BPS Guide to Pharmacology (GtoPdb) [1]. Using a curated dataset of 252 known endogenous ligand–receptor pairs covering 120 GPCRs and 114 ligands, we evaluated the ability of G-LEAP to rediscover established biology by ranking ∼217,000 HMDB metabolites for each receptor. G-LEAP ranked the true endogenous ligand within the top 1% of all candidates for 33.7% of pairs and within the top 5% for 47.2%, demonstrating enrichment far exceeding random expectation **(Fig. 4B)**. Given that random selection would yield a recovery rate of only 1%, this represents a ∼34-fold enrichment over random expectation. This result is particularly striking because G-LEAP was trained primarily on synthetic, drug-like molecules from the GLASS database. Its ability to systematically pinpoint endogenous metabolites suggests that the model has successfully learned transferable principles of molecular recognition from synthetic pharmacology to natural physiology.

We also evaluated the model’s target fishing capability by predicting receptor targets for a given metabolite. For promiscuous ligands such as serotonin, dopamine, and acetylcholine, 78% of their known GPCR partners ranked within the top 10% of 624 GPCRs (mean percentile: 7.1%; **Supplementary Fig. 7B**). This bidirectional validation confirms that G-LEAP captures genuine molecular recognition principles rather than dataset-specific biases.

To further investigate the mechanism of ligand identification, we analyzed the chemical structures of false positives (FP), which are compounds ranked in the top 100 but not annotated as ground truths. Globally, these candidates showed a low mean Tanimoto similarity of 0.111 to the true ligands **(Supplementary Fig. 7C)**, indicating that G-LEAP generally does not rely on simple structural resemblance. Instead, the model captures deeper compatibility signals such as protein-sequence patterns and learned interaction motifs. However, when we analyzed FPs separately for each GPCR family, a clear exception emerged. For Sphingosine-1-phosphate receptors (S1P1, S1P3, and S1P5), over 50% of the FPs exhibited moderate-to-high structural similarity (Tanimoto > 0.3) to the true ligand. A similar trend was observed for the adenosine A3 receptor **(Supplementary Fig. 7D)**. These results indicate that, for these receptors, G-LEAP preferentially assigns high scores to compounds sharing substructural features with the cognate endogenous ligand, consistent with the model implicitly capturing ligand-class-specific recognition patterns. This capacity for fine-grained molecular discrimination was further confirmed by the high-confidence predictions for well-characterized pairs, such as 5-HT receptors with serotonin and MTNR1A with melatonin, which consistently showed high prediction scores above 0.95 and low uncertainty (CV-SD < 0.05), indicating that the model’s confidence is stable across folds rather than driven by a single favorable split **(Fig. 4C)**.

The model’s sensitivity to fine-grained chemical details was exemplified by GPR32, for which Resolvin D1 has been proposed as a candidate endogenous ligand [49]. G-LEAP not only correctly identified Resolvin D1 with a high prediction score but also enriched its structural isomers and analogs (e.g., Resolvin D5, 15-Epi-lipoxin B5) in the top ranks, clearly distinguishing them from other lipid mediators **(Fig. 4D)**. The high structural similarity between the predicted top-scoring compounds and Resolvin D1 confirms that the model extracts data-driven pharmacophore-like features that align with structural and stereochemical constraints.

Finally, applying this validated framework to 23 Class A orphan GPCRs yielded high-confidence predictions that align with phylogenetic and functional insights **(Fig. 4E)**. For GPR26, which is a member of the SREB (super-conserved receptor expressed in brain) family [50], the model predicted the amine-derived melatonin with a score of 0.978 ± 0.018. For GPR20, phylogenetically related to lysolipid receptors [51], the model identified sphingosine 1-phosphate and lysophosphatidylethanolamine (LysoPE). For GPR176 [52], it identified the endocannabinoid anandamide with a score of 0.959 ± 0.026. To assess physiological plausibility, we examined tissue co-expression patterns using GTEx data [53], testing whether the biosynthetic enzymes responsible for producing the predicted endogenous ligands are co-expressed with their cognate orphan receptors across human tissues. Predicted receptor–enzyme pairs showed significant positive correlations, including GPR20 with sphingosine kinase 1 (SPHK1, ρ = 0.627, p < 10^-8^) [54], and GPR26 with arylalkylamine *N*-acetyltransferase (AANAT, ρ = 0.339, p < 0.01) [55] **(Supplementary Fig. 8A, B)**. These findings demonstrate that G-LEAP captures not only binding potential but also the physiological context of ligand candidates, offering a systematic roadmap for accelerating the experimental de-orphanization of the GPCRome.

## Discussion

The de-orphanization of G protein-coupled receptors (GPCRs) remains one of the most formidable challenges in modern pharmacology. While high-throughput screening and structural biology have mapped the “lighted” regions of the GPCRome, hundreds of orphan receptors remain in the “dark matter,” largely inaccessible due to the lack of known ligands or high-resolution structures. This study presents G-LEAP, a geometric-evolutionary deep learning framework that effectively bridges this knowledge gap. By achieving high generalization performance on the rigorous Protein split benchmark, G-LEAP demonstrates a unique capability to predict ligand interactions for unseen GPCRs. Our results suggest that the model has learned transferable principles of molecular recognition that generalize across the GPCR superfamily, enabling predictions for orphan receptors without requiring target-specific structural information.

From a computational perspective, the key innovation of G-LEAP lies in the synergistic integration of evolutionary and geometric modalities. Traditional methods have largely treated protein sequences and ligand structures as separate, static feature sets. In contrast, our dual-encoder architecture leverages the evolutionary context embedded in protein language models and the stereochemical awareness of geometric deep learning. Rather than relying on simple concatenation, we employ a bilinear interaction module that models pairwise compatibility between protein and ligand latent spaces, capturing nonlinear interactions that would be lost in linear feature fusion. This design enables the model to discriminate active ligands based on structural complementarity rather than superficial physicochemical similarity, as demonstrated by its ability to distinguish true binders from property-matched decoys and reject false-positive docking hits. By bypassing the need for manual feature engineering or binding pocket coordinates, G-LEAP enables ligand prediction against structurally uncharacterized targets that would be inaccessible to structure-based methods.

Biologically, G-LEAP serves as a validated discovery engine that translates computational predictions into testable physiological hypotheses. The construction of the GPCR-Metabolome Atlas represents a significant resource for the community, providing a systematic roadmap for experimental de-orphanization efforts. Importantly, our predictions are supported by multi-layered validation: ∼34-fold enrichment over random expectation in recovering known endogenous ligands (Top 1% recall = 33.7% across 252 IUPHAR/BPS-annotated pairs), and significant tissue-specific co-expression of predicted orphan receptor–ligand pairs (e.g., GPR20–sphingosine 1-phosphate) with their biosynthetic enzymes across human tissues. These independent lines of evidence demonstrate that G-LEAP captures not only binding potential but also physiological plausibility. In the context of drug discovery, G-LEAP demonstrated robust scaffold-hopping capacity, identifying structurally novel hits for therapeutic targets that are chemically distinct from existing drugs. This capability is particularly valuable for accessing underexplored regions of chemical space and circumventing the intellectual property constraints associated with traditional derivative-based drug design.

Despite these advances, several limitations delineate the path for future research. First, G-LEAP currently operates on static representations and does not explicitly account for the dynamic conformational landscapes of GPCRs or the lipid membrane environment. Integrating data from molecular dynamics simulations or distinct conformational states could refine predictions and potentially enable the discrimination of functional efficacy, such as agonist versus antagonist activity. Second, while we employed rigorous cross-validation strategies to mitigate dataset biases, the training data remain skewed toward Class A receptors, which constitute 96.0% of available GPCR–ligand interaction data. Strategies such as active learning, few-shot transfer learning, or incorporation of broader interaction databases could further enhance performance for under-represented families including Class B, C, and adhesion GPCRs (Class B2). Finally, while our method predicts binding probability with high confidence, experimental validation remains essential for confirming biological activity, functional selectivity, and downstream signaling consequences.

In conclusion, G-LEAP represents a foundational step toward predictive pharmacology at the GPCRome scale. By successfully generalizing GPCR–ligand recognition patterns without relying on target-specific priors, it provides a versatile platform applicable to dissecting basic physiology and designing next-generation therapeutics. As the volume of genomic and chemical data continues to expand, frameworks like G-LEAP will be essential for transforming this information into actionable biomedical insights, ultimately illuminating the unexplored frontiers of the human GPCRome and accelerating the discovery pipeline for orphan receptor pharmacology.

## Materials and Methods

### Data collection and curation

To construct a comprehensive dataset for GPCR–ligand interaction prediction, we integrated data from three primary repositories: GLASS [14], GPCRdb [15], and the recently released GLASS 2.0 [21].

### Initial dataset construction (GLASS and GPCRdb)

For the initial model development phase, we aggregated interaction records from GLASS [14] and GPCRdb [15]. From GLASS, we extracted 1,045,681 interaction records containing quantitative affinity metrics (IC_50_, K_i_, or K_d_). Chemical structures were retrieved as SMILES (Simplified Molecular Input Line Entry System) strings from the associated SDF (structure-data file) files. From GPCRdb, we retrieved 403,009 interaction pairs for human GPCRs via the REST API.

Chemical structures were standardized using the RDKit library (version 2023.09.5) [18] in Python. The preprocessing pipeline included: (1) conversion to canonical SMILES, (2) removal of salts and solvents, (3) normalization of functional groups, (4) standardization of protonation states, and (5) tautomer canonicalization.

Using the standardized SMILES and UniProt IDs as unique identifiers, we merged the two databases and removed duplicate entries. For pairs with multiple recorded affinity values (K_i_, pIC_50_), the mean value was calculated. We applied strict bioactivity thresholds to define binary labels: pairs with an affinity < 4 nM (pIC_50_ > 8.40) were labeled as active (label=1), and those with an affinity > 4,000 nM (pIC_50_ < 5.40) were labeled as inactive (label=0), as previously reported [10]. This process yielded an initial dataset of 120,291 unique pairs (612 proteins, 81,679 compounds) with a balanced distribution (56,574 actives, 63,717 inactives).

### Benchmark dataset construction (GLASS 2.0)

To evaluate the model on the most up-to-date and extensive data available, we processed the GLASS 2.0 database [21]. We combined 482,645 active records and 81,742 inactive records. Records with non-numeric or missing standard values were excluded. To ensure a rigorous binary classification benchmark, we adopted a stringent gap-based thresholding strategy. Pairs with an affinity ≤ 10 nM were labeled as positive, while pairs with an affinity ≥ 1,000 nM were labeled as negative. Pairs falling into the intermediate “gray zone” (10 nM < affinity < 1,000 nM) were excluded to remove ambiguous data points that could hinder model convergence. We resolved duplicate entries (pairs with identical UniProt ID and canonical SMILES) using a majority vote strategy: the most frequent label was assigned as the representative class. In cases of a tie (equal frequency of positive and negative labels), the pair was conservatively labeled as negative. Following these filtration and de-duplication steps, the final GLASS 2.0 benchmark dataset consisted of 291,679 unique pairs (114,644 positives; 177,035 negatives), covering 854 unique GPCRs and 180,819 unique compounds.

### Data splitting strategies

To rigorously assess the model’s generalization capabilities across different domains, we implemented three distinct data splitting strategies. For the initial dataset, we partitioned the data into training, validation, and test sets with an 8:1:1 ratio. First, a “Random split” was performed using stratified sampling to maintain the ratio of positive to negative samples, serving as a baseline evaluation. Second, a “Ligand split” was implemented based on unique canonical SMILES, ensuring that compounds in the test set were entirely absent from the training set; this evaluates the model’s ability to generalize to novel chemical scaffolds. Third, a “Protein split” was constructed based on unique UniProt IDs, ensuring that receptors in the test set were unseen during training. This split is critical for simulating the de-orphanization scenario, evaluating the model’s capacity to predict ligands for novel targets. For the GLASS 2.0 dataset, we adopted a 10-fold cross-validation (CV) scheme to ensure statistical robustness. Like the initial strategy, we employed 10-fold Random, Ligand, and Protein splits. In the Ligand and Protein splits, the unique compounds or receptors were randomly partitioned into 10 folds, ensuring that each test fold consisted entirely of unseen chemical or biological entities, respectively. These split datasets were also utilized to retrain and evaluate baseline methods (BIND [22], DEAttentionDTA [23], and Perceiver CPI [24]) to ensure fair comparison. All data processing and splitting procedures were implemented using Python (Pandas v2.2.3, NumPy v1.26.4, scikit-learn v1.5.2).

## Feature preparation

### Compound feature engineering and graph construction

Input compounds were represented as molecular graphs comprising atom-level node features and bond-based edges. Molecular graphs were constructed from SMILES strings using RDKit (version 2024.09.1), where heavy atoms (excluding hydrogens) served as nodes and chemical bonds served as edges; no distinction was made regarding bond orders (e.g., single or double bonds). To capture rich chemical information, we employed a hybrid node featurization strategy combining deep learning embeddings with physicochemical descriptors. For the deep learning component, we utilized Uni-Mol2 [20], an advanced molecular pre-training framework that integrates atom-level, graph-level, and geometric features. We employed the pre-trained Uni-Mol2 model (164 million parameters) to extract a 768-dimensional embedding for each atom. Complementing this, we generated 113-dimensional handcrafted feature vectors using RDKit, which encoded physicochemical properties including atom type, degree, number of bonded hydrogens, implicit valence, aromaticity, formal charge, chirality, ring membership, and hydrogen bond donor/acceptor status. These two feature sets were concatenated to yield a final 881-dimensional vector for each atomic node, which served as the input for the G-LEAP framework. For benchmarking experiments against prior art, we also generated features according to the DeepGPCR protocol [10]. For compounds, atoms were represented by 78-dimensional vectors constructed via one-hot encoding of five attributes: atom type (44 categories), degree (0–10), number of bonded hydrogens (0–10), implicit valence (0–10), and aromaticity. For proteins, each of the 20 standard amino acids was represented by a 30-dimensional embedding generated via the mol2vec algorithm [17].

### Protein structural representation and evolutionary embedding

Protein inputs were similarly processed into graph structures derived from 3D conformational data. Structural coordinates for human GPCRs were primarily retrieved from the AlphaFold Protein Structure Database (AlphaFold DB; downloaded October 17, 2024) [56]. For receptors not available in the database, 3D structures were individually predicted using AlphaFold2 [57]. We constructed residue-level graphs where each amino acid residue represented a node centered on its α-carbon (C_α_). Unweighted edges were defined between residue pairs if the Euclidean distance between their C_α_ atoms was less than 5 Å, capturing local structural neighborhoods. Node features were extracted using the protein language model ESM Cambrian (ESM C) [19], which captures deep evolutionary constraints from protein sequences. We utilized the open-source ESM C model (300 million parameters) to generate a 960-dimensional embedding vector for each amino acid residue. These evolutionary embeddings served as the node features for the protein graph in the G-LEAP framework.

## Model architecture

### Systematic evaluation of graph neural network backbones

To identify the optimal inductive bias for modeling GPCR–ligand interactions, we evaluated five Graph Neural Network (GNN) architectures based on Graph Attention Networks (GAT) [58], Graph Convolutional Networks (GCN) [16], and Graph Isomorphism Networks (GIN) [59]. We tested both homogeneous (GAT-Bilinear, GCN-Bilinear, GIN-Bilinear) and hybrid (GAT-GCN-Bilinear, GIN-GCN-Bilinear) configurations. All variants shared a common pipeline: protein and ligand graphs were processed by independent encoder streams to extract latent representations, which were then aggregated via global pooling, fused by a shared bilinear interaction module, and passed to a final predictor to output the probability of binding.

### Dual-stream graph encoders

For both ligand and protein streams, we employed a two-layer GNN encoder structure, though the specific operator composition varied by model variant. In GAT-based layers, we utilized graph attention convolutions with 4 attention heads and Exponential Linear Unit (ELU) activation to capture anisotropic neighborhood features. GCN and GIN layers employed spectral graph convolutions and isomorphism operators, respectively, both utilizing Rectified Linear Unit (ReLU) activation. Hybrid architectures leveraged complementary strengths; for instance, the GAT-GCN variant consisted of a first-layer GATConv followed by a second-layer GCNConv. In all configurations, dropout regularization was applied between layers to prevent overfitting. Following the graph convolutions, node-level feature vectors were aggregated into a fixed-size graph-level representation using global mean pooling. While the input dimensions corresponded to the respective feature sets (d_ligand_=881, d_protein_=960), all encoders projected these inputs into a unified latent embedding dimension (embed_dim) to facilitate downstream integration.

### Bilinear interaction module

Given the pooled ligand and protein embeddings *x_l_* and *x_p_*, we computed an interaction vector *Z* ∈ ℝ*^H^* using a bilinear transformation implemented with PyTorch’s nn.Bilinear (input dimensions = embed_dim, output dimension = H). The vector *Z* was passed through ReLU and dropout, and a scalar gating coefficient α ∈ [0,1] was produced via a multilayer perceptron (MLP) followed by a sigmoid. The gated interaction α ⋅ *Z* was then transformed by an additional MLP, yielding the final fused representation.

### Final predictor

The fused representation was fed into a fully connected network with ReLU activations and dropout. The final sigmoid output produced the predicted probability that a given GPCR–ligand pair is active.

## Model training and evaluation

### Training Protocol

The framework was trained as a binary classification task to distinguish between active and inactive GPCR–ligand pairs. We optimized the model parameters using the Adam optimizer with a learning rate of 1.0×10^-3^ and a weight decay of 1.0×10^-5^ to enforce regularization. The binary cross-entropy loss function was minimized over a maximum of 100 epochs using a batch size of 128. The architecture was configured with a unified embedding dimension of 128 and a hidden dimension of 256 for the bilinear block, with a dropout rate of 0.2 applied across network layers to mitigate overfitting. To ensure optimal model convergence and prevent overtraining, we implemented an early stopping mechanism with a patience of 20 epochs, which automatically terminated training if the validation loss failed to improve. Independent training sessions were conducted for each of the three data splitting strategies (Random, Ligand, and Protein splits) to ensure a comprehensive assessment of the model’s learning capabilities under different generalization scenarios.

### Evaluation metrics

Model performance was rigorously evaluated on the held-out test sets using a comprehensive suite of statistical metrics. We primarily utilized threshold-independent metrics—specifically the Area Under the Receiver Operating Characteristic Curve (ROC-AUC) and the Average Precision (AP) —to assess the global discriminative power of the model across all decision thresholds. Additionally, to provide a tangible measure of classification performance, we calculated threshold-dependent metrics including Accuracy, Precision, Recall, and F1-score. For these binary metrics, a standard decision threshold of 0.5 was applied to the predicted probability scores.

### Hyperparameter tuning

We conducted a comprehensive hyperparameter optimization using a constrained grid search strategy. A total of 97 candidate configurations were evaluated across learning rate (0.0005, 0.0008, 0.001, 0.0012), weight decay (5×10⁻⁶, 1×10⁻⁵, 2×10⁻⁵, 5×10⁻⁵), batch size (64, 96, 128, 160, 192), embedding dimension (96, 128, 160, 192, 224, 256), dropout rate (0.15, 0.2, 0.25, 0.3), optimizer (Adam, AdamW), scheduler (cosine, plateau), and normalization (none, batch). All configurations were initially screened using fold 0 of the protein-based cross-validation split. The top 10 configurations were then subjected to full 10-fold cross-validation to ensure statistically reliable performance evaluation. The final optimal hyperparameters were determined based on 10-fold CV mean performance: learning rate = 0.001, weight decay = 1×10⁻⁵, batch size = 128, embedding dimension = 128, dropout = 0.2, Adam optimizer, plateau scheduler, and batch normalization (10-fold CV ROC-AUC = 0.829 ± 0.028).

### GPCR Family-Wise Performance Evaluation

To assess the robustness of the model across the diverse GPCR superfamily, we performed a stratified performance evaluation using the held-out test sets derived from the 10-fold cross-validation procedure. Receptor class annotations were assigned by mapping UniProt IDs to their respective families (Class A, B, C, F, and others) using the hierarchical classification scheme provided by GPCRdb [15]. Of the 287,653 total prediction instances, 188,823 entries (65.6%) were successfully mapped to specific family annotations. Performance metrics—including Accuracy, ROC-AUC, AP, and F1-score—were computed for each GPCR family by aggregating predictions across all 10 cross-validation folds. Given the pronounced class imbalance inherent in the annotated dataset—which was heavily dominated by Class A (96.0%), followed by Class C (2.8%), Class B (1.0%), and Class F (0.1%)—we reported both threshold-independent metrics (ROC-AUC, AP) to assess global ranking capability and threshold-dependent metrics (Accuracy, F1-score) calculated at a default decision boundary of 0.5.

### Benchmark Datasets

#### DUDE-Z

To evaluate the screening performance of G-LEAP, we used a benchmark dataset comprising known ligands (actives) and non-binders (decoys) with closely matched physicochemical properties. In this study, we employed the DUDE-Z database [25], which includes particularly challenging decoys (accessed December 24, 2025). From DUDE-Z, we retrieved known ligands and decoys for three GPCR targets: dopamine receptor D4 (DRD4; UniProt: P21917), adenosine receptor A2a (ADORA2A; UniProt: P29274), and melatonin receptor type 1A (MTNR1A; UniProt: P48039). For each GPCR, compounds were explicitly labeled as “ligand” (known ligands) or “decoy” (decoys), yielding an evaluation-ready dataset that can be readily compared with downstream inference results. In total, we obtained DRD4 (ligand:decoy = 37:1850), ADORA2A (90:4500), and MTNR1A (28:2150) sets.

#### External validation dataset (GPR139)

To assess the generalizability of our model, we used an independent external validation set from Cabeza de Vaca *et al.*, who reported the discovery of GPR139 agonists through ultra-large virtual screening [28]. From this study, we extracted 68 compounds tested in the initial virtual screening iteration. Of these, five compounds showed agonist activity in Ca²⁺ mobilization assays with EC₅₀ values ranging from 160 nM to 3.6 µM (pEC₅₀: 5.4–6.8), while the remaining 63 compounds were inactive. This dataset provides an orthogonal benchmark, as these compounds were identified using physics-based docking (DOCK3.7) against the GPR139 cryo-EM structure (PDB: 7VUG), representing a methodologically distinct approach from our AI-based prediction.

### Virtual library preparation

To explore the chemical space for both endogenous substrates and novel therapeutic candidates, we prepared two distinct screening libraries.

#### Human Metabolome Database (HMDB)

For the systematic de-orphanization of GPCRs, we retrieved metabolite structures from the Human Metabolome Database [47] (accessed July 25, 2025). SDF files were converted to canonical SMILES strings using RDKit. These compounds were subjected to the same preprocessing and feature engineering pipeline as the training data, resulting in a library of approximately 217,000 endogenous metabolites.

#### ZINC20 Dataset

For large-scale *in silico* drug screening, we sourced Reactive Standard and Purchasable compounds from the ZINC20 database (accessed August 1, 2025) [31]. To ensure the pharmacological viability of the candidates, we applied a rigorous medicinal chemistry filtration cascade. First, we filtered compounds based on Lipinski’s rule of five (H-bond donors ≤ 5, H-bond acceptors ≤ 10, LogP ≤ 5, Molecular Weight ≤ 500) to ensure oral bioavailability [37]. Second, we removed Pan-Assay Interference Compounds (PAINS) to exclude promiscuous binders and frequent hitters [38]. Third, we applied Brenk’s Structural Alerts to filter out compounds with potential toxicity or reactive instability [39]. Following these filtration steps, the remaining compounds were processed into feature vectors identical to the training protocol, yielding a high-quality screening library of 6,151,237 drug-like compounds.

#### GPCR targets

Virtual screening was performed against the following GPCRs: free fatty acid receptor 1 (FFAR1, UniProt: O14842), the muscarinic acetylcholine receptor M2 (CHRM2, UniProt: P08172), the orexin receptor type 2 (HCRTR2, UniProt: O43614), GPR3 (UniProt: P46089), GPR17 (UniProt: Q13304), and GPR88 (UniProt: Q9GZN0).

### Fingerprint-based similarity thresholds for novelty assessment

To compare structural similarity between compounds, we established fingerprint-specific reference thresholds (novelty cutoffs) using multiple molecular fingerprints. We used three commonly employed fingerprints: Morgan fingerprint [60], RDKit fingerprint, and MACCS keys [61]. Compounds were converted to bit vectors with RDKit (version 2024.9.6) using rdkit.Chem.AllChem.GetMorganFingerprintAsBitVect for Morgan fingerprints, rdkit.Chem.rdFingerprintGenerator.GetRDKitFPGenerator for RDKit fingerprints, and rdkit.Chem.MACCSkeys.GenMACCSKeys for MACCS keys. Parameters were set to radius = 2 and nBits = 1024 for Morgan fingerprints, and fpsize = 2048 and maxPath = 7 for RDKit fingerprints; all other parameters were kept at their default values. To derive reference thresholds, we randomly retrieved 1,000 compounds from PubChem via its API (accessed January 7, 2026) and computed pairwise Tanimoto similarities for all unique pairs. We then analyzed the resulting similarity distributions and defined, for each fingerprint, a novelty cutoff corresponding to high similarity values that are rarely observed among random compound pairs.

### Enrichment factor analysis

To evaluate whether G-LEAP can discriminate ligands from decoys, we computed the enrichment factor (EF) for each GPCR. All compounds (ligand and decoy) were ranked by their G-LEAP scores, and the top X% was defined based on this global ranking. EF at X% (*EF*_*X*%) was calculated as:

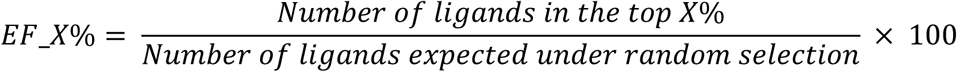

We reported EF as X = [0.1, 0.2, 0.5, 1, 2, 5, 10] in this study.

GPCRs used for EF analysis were D4 dopamine receptor (DRD4, UniProt: P21917), the adenosine receptor A2a (ADORA2A, UniProt: P29274), and the melatonin receptor type 1A (MTNR1A, UniProt: P48039).

### Chemical novelty analysis relative to known ligands

To assess the novelty of compounds identified by G-LEAP, we established a method to quantify the structural distance of each compound relative to the set of known ligands. For each target GPCR *t*, we collected a reference set of known ligands *L_t_* and a set of G-LEAP high-scoring compounds *S*. To ensure that novelty estimates were not dependent on a single molecular representation, each compound was encoded in three commonly used fingerprint types (Morgan, MACCS keys, and RDKit fingerprints). For each high-scoring compound *s* ∈ *S* and target GPCR *t*, we computed the maximum similarity to any known ligand in *L_t_*:

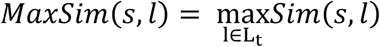

Similarity, *Sim*(*s*, *l*), was computed using the Tanimoto coefficient. This procedure identifies, for each predicted hit, its nearest neighbor among known ligands in the chosen fingerprint space.

Based on *Sim*(*s*, *l*), we computed pairwise similarities between compounds randomly retrieved from PubChem via the PubChem API (accessed January 7, 2026).

GPCRs used for chemical novelty analysis were free fatty acid receptor 1 (FFAR1, UniProt: O14842), the muscarinic acetylcholine receptor M2 (CHRM2, UniProt: P08172), and the orexin receptor type 2 (HCRTR2, UniProt: O43614).

### Chemical space mapping for novelty assessment

To visualize the novelty of compounds identified by G-LEAP, we computed Morgan fingerprints (radius = 2, nBits = 2048) using RDKit (version: 2024.9.6) and performed dimensionality reduction by UMAP (n_neighbors = 15, min_dist = 0.1, metric = “jaccard”; version: 0.5.9) within Python (version 3.10.17). The UMAP embedding was fitted using all compounds included in the plot, and the resulting coordinates were used for visualization.

### Structural modeling and affinity estimation via Boltz-2

To validate the physical plausibility of the G-LEAP predictions, we employed Boltz-2 [40] for 3D structural modeling and binding affinity estimation of selected GPCR–ligand complexes. Multiple sequence alignments (MSA) required for protein folding were generated via the –-use_msa_server option. To balance computational efficiency with prediction accuracy for the large-scale validation, we configured the sampling parameters as follows: diffusion_sample = 5, recycling_step = 5, and max_parallel_sample = 5; all other parameters were maintained at default settings.

Binding affinities were derived from the model’s confidence outputs. Following the official Boltz-2 documentation, the predicted affinity value (affinity_pred_value) was converted to an estimated IC_50_ using the following equation:

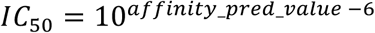

This physics-informed validation step served to corroborate the deep learning predictions with structural evidence of binding stability.

### Molecular dynamics (MD) simulation and preprocessing

MD simulations were performed to evaluate the binding stability of the top-ranked compounds identified through G-LEAP screening with the target GPCRs. The Molecular Operating Environment (MOE, Chemical Computing Group Inc.) [62] was used for MD simulations. Initially, the complex structure models were prepared to complete missing atoms, add hydrogen atoms, assign tether restraints, and perform structural optimization using the Amber10:ETH force field [63]. The initial structures were taken from the Boltz-2–predicted models and the optimized structures were then utilized for both docking calculations and MD simulations.

### Docking calculations

Docking calculations were performed using MOE [62]. Initially, the stability of the compound on the protein surface was evaluated using simulation with a time step of 1 fs. Once the energy had converged, the binding free energy between the protein and the compound was calculated. Fifty initial poses of the compound were generated using the Triangle Matcher placement algorithm, and the calculations were repeated for each pose. The binding free energy was estimated using the London ΔG scoring function [64], as shown in the following equation:

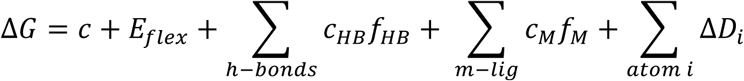

Where *c* represents the average loss of rotational and translational entropy of the compound, *E_flex_* is the energy loss due to ligand flexibility calculated solely from its conformation, *f_HB_* is a variable (ranging from 0 to 1) representing the geometric imperfection of hydrogen bonds, *C_HB_* is the ideal hydrogen bond energy, *f_M_* is a variable (ranging from 0 to 1) representing the geometric imperfection of metal-ligand interactions, *c_M_* is the ideal energy for metal-ligand interactions, and *D*_*i*_ represents the desolvation energy of the *i*th atom of the ligand, which is calculated as follows:

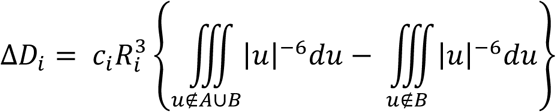

In this equation, *A* and *B* denote the spatial regions occupied by the protein and ligand, respectively, *R_i_* is the solvation radius of the *i* th atom, and *c*_*i*_ is a constant specific to the atom type.

Additionally, the binding free energy was calculated using the GBVI/WSA ΔG function [65], according to the following expression:

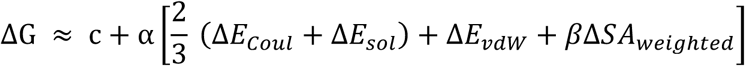

where *c* represents the average loss of rotational and translational entropy of the compound, **E*_coul_* is the electrostatic energy, *E_sol_* is the solvation electrostatic energy, *E_vdw_* is the van der Waals energy, *SA_weighted_* is the weighted solvent-accessible surface area, and α and β are constants.

In this study, 30 initial poses were generated for each compound, and up to 10 poses ranked by predicted binding free energy (ΔG) were retained. The reported binding energy (kcal/mol) corresponds to the minimum value among the retained poses. Protein-ligand interaction fingerprints (PLIFs) were calculated using the set of retained poses for each compound.

## Statistical analysis

Statistical analysis was performed using the scipy.stats module (version 1.15.2) within Python (version 3.10.17). Comparison of binding energy between high-scoring group and low-scoring group was assessed using a two-sided Mann–Whitney *U-*test. A *p*-value less than 0.05 was considered statistically significant.

## ACKNOWLEDGEMENTS

This work was supported by KAKENHI grants from the Japan Society for the Promotion of Science (JSPS) to H.S. (23K28184, 24H01755 and 25H01571), JST FOREST Program to H.S. (JPMJFR242Q), as well as the Canon Foundation and Nakatani Foundation. Figures were created in part with BioRender.com. We thank T. Suzuoka, T. Suzuki, K. Nakanishi, and T. Ito for critical reading of the manuscript, all laboratory members for helpful discussions, and K. Tanaka for assistance with manuscript preparation.

## AUTHOR CONTRIBUTIONS

T.S. initially conceived of the project. H.S. designed and supervised the study.

T.S. and Y.O. developed G-LEAP and performed all formal analyses. T.S., Y.O. and H.S. jointly wrote the manuscript. All authors have read and approved the final manuscript.

