## Supplementary Tables and Figures for "Geometric-Evolutionary Deep Learning Decodes the Human GPCR-Metabolome Interactome and Enables Systematic De-Orphanization"

**Supplementary Table 1 | Comparison of GNN backbone architectures with bilinear interaction module on the GLASS dataset**

| GNN Architecture | Random (ROC-AUC) | Ligand (ROC-AUC) | Protein (ROC-AUC) |
|---|---|---|---|
| GCN-Bilinear | 0.984 | 0.985 | 0.890 |
| GAT-Bilinear | 0.984 | 0.982 | 0.871 |
| GAT-GCN-Bilinear | 0.985 | 0.985 | 0.859 |
| GIN-Bilinear | 0.965 | 0.685 | 0.823 |
| GIN-GCN-Bilinear | 0.500 | 0.511 | 0.501 |

GCN, Graph Convolutional Network; GAT, Graph Attention Network; GIN, Graph Isomorphism Network. GAT-GCN and GIN-GCN denote hybrid architectures using GAT or GIN for ligand graphs and GCN for protein graphs, respectively.

**Supplementary Table 2 | Hyperparameter configurations and ROC-AUC performance from initial fold 0 screening in G-LEAP optimization**

| Rank | AUC | LR | WD | Batch | Embed Dim | Dropout | Layers | Optimizer | Scheduler | Norm | Pool |
|---|---|---|---|---|---|---|---|---|---|---|---|
| 1 | 0.8348 | 0.0012 | 1e-05 | 128 | 224 | 0.20 | 2 | adamw | cosine | none | mean |
| 2 | 0.8336 | 0.0012 | 1e-05 | 64 | 128 | 0.20 | 2 | adamw | cosine | none | mean |
| 3 | 0.8333 | 0.0008 | 1e-05 | 128 | 128 | 0.20 | 2 | adam | cosine | none | mean |
| 4 | 0.8322 | 0.0010 | 1e-05 | 128 | 128 | 0.20 | 2 | adam | plateau | batch | mean |
| 5 | 0.8313 | 0.0010 | 1e-05 | 160 | 128 | 0.20 | 2 | adamw | cosine | none | mean |
| 6 | 0.8310 | 0.0010 | 5e-05 | 128 | 96 | 0.20 | 2 | adamw | cosine | none | mean |
| 7 | 0.8308 | 0.0010 | 5e-05 | 128 | 192 | 0.20 | 2 | adamw | cosine | none | mean |
| 8 | 0.8302 | 0.0008 | 1e-05 | 128 | 256 | 0.20 | 2 | adamw | cosine | none | mean |

| Rank | AUC | LR | WD | Batch | Embed Dim | Dropout | Layers | Optimizer | Scheduler | Norm | Pool |
|---|---|---|---|---|---|---|---|---|---|---|---|
| 9 | 0.8299 | 0.0008 | 2e-05 | 128 | 192 | 0.20 | 2 | adamw | cosine | none | mean |
| 10 | 0.8299 | 0.0010 | 2e-05 | 128 | 128 | 0.20 | 2 | adam | plateau | none | mean |
| 11 | 0.8290 | 0.0008 | 1e-05 | 128 | 96 | 0.20 | 2 | adamw | cosine | none | mean |
| 12 | 0.8282 | 0.0010 | 1e-05 | 128 | 192 | 0.15 | 2 | adamw | cosine | none | mean |
| 13 | 0.8273 | 0.0010 | 1e-05 | 128 | 160 | 0.30 | 2 | adam | plateau | none | mean |
| 14 | 0.8272 | 0.0008 | 1e-05 | 128 | 160 | 0.20 | 2 | adamw | cosine | none | mean |
| 15 | 0.8271 | 0.0012 | 1e-05 | 128 | 256 | 0.20 | 2 | adamw | cosine | none | mean |
| 16 | 0.8268 | 0.0008 | 2e-05 | 128 | 256 | 0.20 | 2 | adamw | cosine | none | mean |
| 17 | 0.8266 | 0.0008 | 2e-05 | 128 | 128 | 0.20 | 2 | adam | plateau | none | mean |
| 18 | 0.8262 | 0.0008 | 2e-05 | 128 | 96 | 0.20 | 2 | adamw | cosine | none | mean |
| 19 | 0.8262 | 0.0010 | 1e-05 | 192 | 128 | 0.20 | 2 | adamw | cosine | none | mean |
| 20 | 0.8260 | 0.0008 | 2e-05 | 128 | 160 | 0.20 | 2 | adamw | cosine | none | mean |
| 21 | 0.8256 | 0.0008 | 1e-05 | 96 | 128 | 0.20 | 2 | adamw | cosine | none | mean |
| 22 | 0.8249 | 0.0010 | 1e-05 | 128 | 128 | 0.20 | 2 | adamw | plateau | none | mean |
| 23 | 0.8248 | 0.0010 | 1e-05 | 128 | 256 | 0.20 | 2 | adam | plateau | none | mean |
| 24 | 0.8246 | 0.0010 | 5e-06 | 128 | 128 | 0.20 | 2 | adam | plateau | none | mean |
| 25 | 0.8245 | 0.0010 | 1e-05 | 128 | 160 | 0.15 | 2 | adam | plateau | none | mean |
| 26 | 0.8239 | 0.0008 | 2e-05 | 128 | 224 | 0.20 | 2 | adamw | cosine | none | mean |
| 27 | 0.8238 | 0.0012 | 2e-05 | 128 | 160 | 0.20 | 2 | adamw | cosine | none | mean |
| 28 | 0.8237 | 0.0012 | 1e-05 | 96 | 128 | 0.20 | 2 | adamw | cosine | none | mean |
| 29 | 0.8236 | 0.0012 | 1e-05 | 128 | 160 | 0.20 | 2 | adamw | cosine | none | mean |
| 30 | 0.8221 | 0.0010 | 2e-05 | 128 | 224 | 0.20 | 2 | adamw | cosine | none | mean |
| 31 | 0.8217 | 0.0010 | 5e-06 | 128 | 224 | 0.20 | 2 | adamw | cosine | none | mean |
| 32 | 0.8211 | 0.0010 | 1e-05 | 128 | 128 | 0.20 | 2 | adam | plateau | none | mean |
| 33 | 0.8209 | 0.0010 | 1e-05 | 64 | 128 | 0.20 | 2 | adam | plateau | none | mean |
| 34 | 0.8205 | 0.0012 | 1e-05 | 160 | 128 | 0.20 | 2 | adamw | cosine | none | mean |
| 35 | 0.8200 | 0.0010 | 1e-05 | 160 | 128 | 0.20 | 2 | adam | plateau | none | mean |
| 36 | 0.8193 | 0.0008 | 1e-05 | 128 | 128 | 0.20 | 2 | adam | plateau | none | mean |
| 37 | 0.8180 | 0.0012 | 5e-06 | 128 | 128 | 0.20 | 2 | adam | plateau | none | mean |
| 38 | 0.8179 | 0.0010 | 1e-05 | 128 | 224 | 0.20 | 2 | adamw | cosine | none | mean |

| Rank | AUC | LR | WD | Batch | Embed Dim | Dropout | Layers | Optimizer | Scheduler | Norm | Pool |
|---|---|---|---|---|---|---|---|---|---|---|---|
| 39 | 0.8172 | 0.0010 | 1e-05 | 96 | 128 | 0.20 | 2 | adamw | cosine | none | mean |
| 40 | 0.8172 | 0.0010 | 1e-05 | 128 | 192 | 0.20 | 2 | adamw | cosine | none | mean |
| 41 | 0.8169 | 0.0010 | 1e-05 | 128 | 128 | 0.20 | 2 | adam | plateau | none | mean |
| 42 | 0.8168 | 0.0010 | 1e-05 | 128 | 160 | 0.15 | 2 | adamw | cosine | none | mean |
| 43 | 0.8166 | 0.0010 | 1e-05 | 128 | 128 | 0.20 | 2 | adam | plateau | none | mean |
| 44 | 0.8159 | 0.0010 | 1e-05 | 128 | 128 | 0.20 | 2 | adam | plateau | none | mean |
| 45 | 0.8157 | 0.0010 | 1e-05 | 64 | 128 | 0.20 | 2 | adamw | cosine | none | mean |
| 46 | 0.8152 | 0.0010 | 5e-06 | 128 | 192 | 0.20 | 2 | adamw | cosine | none | mean |
| 47 | 0.8149 | 0.0008 | 1e-05 | 192 | 128 | 0.20 | 2 | adamw | cosine | none | mean |
| 48 | 0.8145 | 0.0012 | 2e-05 | 128 | 96 | 0.20 | 2 | adamw | cosine | none | mean |
| 49 | 0.8145 | 0.0010 | 1e-05 | 128 | 160 | 0.25 | 2 | adamw | cosine | none | mean |
| 50 | 0.8139 | 0.0010 | 5e-05 | 128 | 160 | 0.20 | 2 | adamw | cosine | none | mean |
| 51 | 0.8138 | 0.0010 | 5e-05 | 128 | 224 | 0.20 | 2 | adamw | cosine | none | mean |
| 52 | 0.8133 | 0.0010 | 5e-06 | 128 | 96 | 0.20 | 2 | adamw | cosine | none | mean |
| 53 | 0.8130 | 0.0010 | 1e-05 | 128 | 224 | 0.20 | 2 | adam | plateau | none | mean |
| 54 | 0.8129 | 0.0010 | 5e-06 | 128 | 256 | 0.20 | 2 | adamw | cosine | none | mean |
| 55 | 0.8126 | 0.0010 | 5e-05 | 128 | 256 | 0.20 | 2 | adamw | cosine | none | mean |
| 56 | 0.8126 | 0.0008 | 1e-05 | 160 | 128 | 0.20 | 2 | adamw | cosine | none | mean |
| 57 | 0.8108 | 0.0010 | 5e-06 | 128 | 160 | 0.20 | 2 | adamw | cosine | none | mean |
| 58 | 0.8106 | 0.0010 | 2e-05 | 128 | 192 | 0.20 | 2 | adamw | cosine | none | mean |
| 59 | 0.8104 | 0.0010 | 1e-05 | 128 | 160 | 0.25 | 2 | adam | plateau | none | mean |
| 60 | 0.8103 | 0.0010 | 1e-05 | 128 | 96 | 0.15 | 2 | adam | plateau | none | mean |
| 61 | 0.8102 | 0.0010 | 1e-05 | 128 | 128 | 0.20 | 2 | adam | plateau | none | mean |
| 62 | 0.8099 | 0.0010 | 1e-05 | 128 | 128 | 0.20 | 2 | adam | plateau | none | mean |
| 63 | 0.8097 | 0.0010 | 1e-05 | 128 | 128 | 0.30 | 2 | adam | plateau | none | mean |
| 64 | 0.8091 | 0.0010 | 5e-05 | 128 | 128 | 0.20 | 2 | adam | plateau | none | mean |
| 65 | 0.8077 | 0.0010 | 2e-05 | 128 | 256 | 0.20 | 2 | adamw | cosine | none | mean |
| 66 | 0.8073 | 0.0012 | 1e-05 | 128 | 192 | 0.20 | 2 | adamw | cosine | none | mean |
| 67 | 0.8071 | 0.0010 | 1e-05 | 128 | 128 | 0.25 | 2 | adam | plateau | none | mean |
| 68 | 0.8070 | 0.0008 | 5e-05 | 128 | 128 | 0.20 | 2 | adam | plateau | none | mean |

| Rank | AUC | LR | WD | Batch | Embed Dim | Dropout | Layers | Optimizer | Scheduler | Norm | Pool |
|---|---|---|---|---|---|---|---|---|---|---|---|
| 69 | 0.8066 | 0.0012 | 5e-05 | 128 | 128 | 0.20 | 2 | adam | plateau | none | mean |
| 70 | 0.8058 | 0.0010 | 1e-05 | 96 | 128 | 0.20 | 2 | adam | plateau | none | mean |
| 71 | 0.8057 | 0.0010 | 1e-05 | 128 | 192 | 0.20 | 2 | adam | plateau | none | mean |
| 72 | 0.8047 | 0.0010 | 1e-05 | 128 | 128 | 0.20 | 2 | adam | plateau | none | mean |
| 73 | 0.8028 | 0.0010 | 1e-05 | 128 | 128 | 0.20 | 4 | adam | plateau | none | mean |
| 74 | 0.8026 | 0.0008 | 5e-06 | 128 | 128 | 0.20 | 2 | adam | plateau | none | mean |
| 75 | 0.8023 | 0.0012 | 1e-05 | 128 | 96 | 0.20 | 2 | adamw | cosine | none | mean |
| 76 | 0.8012 | 0.0008 | 1e-05 | 64 | 128 | 0.20 | 2 | adamw | cosine | none | mean |
| 77 | 0.8010 | 0.0012 | 1e-05 | 192 | 128 | 0.20 | 2 | adamw | cosine | none | mean |
| 78 | 0.8008 | 0.0012 | 1e-05 | 128 | 128 | 0.20 | 3 | adam | cosine | none | mean |
| 79 | 0.8005 | 0.0010 | 1e-05 | 128 | 192 | 0.30 | 2 | adamw | cosine | none | mean |
| 80 | 0.8000 | 0.0010 | 1e-05 | 128 | 128 | 0.20 | 2 | adam | plateau | none | mean |
| 81 | 0.7998 | 0.0010 | 1e-05 | 128 | 256 | 0.20 | 2 | adamw | cosine | none | mean |
| 82 | 0.7997 | 0.0010 | 1e-05 | 128 | 192 | 0.25 | 2 | adamw | cosine | none | mean |
| 83 | 0.7980 | 0.0010 | 1e-05 | 128 | 128 | 0.15 | 2 | adam | plateau | none | mean |
| 84 | 0.7977 | 0.0012 | 1e-05 | 128 | 128 | 0.20 | 2 | adam | plateau | none | mean |
| 85 | 0.7975 | 0.0010 | 1e-05 | 192 | 128 | 0.20 | 2 | adam | plateau | none | mean |
| 86 | 0.7969 | 0.0010 | 1e-05 | 128 | 128 | 0.20 | 2 | adam | plateau | none | mean |
| 87 | 0.7952 | 0.0012 | 2e-05 | 128 | 128 | 0.20 | 2 | adam | plateau | none | mean |
| 88 | 0.7915 | 0.0010 | 1e-05 | 128 | 160 | 0.30 | 2 | adamw | cosine | none | mean |
| 89 | 0.7912 | 0.0010 | 1e-05 | 128 | 128 | 0.20 | 2 | adam | plateau | layer | mean |
| 90 | 0.7863 | 0.0010 | 1e-05 | 128 | 128 | 0.20 | 2 | adam | plateau | batch | add |
| 91 | 0.7855 | 0.0010 | 1e-05 | 128 | 128 | 0.20 | 2 | adam | plateau | none | mean |
| 92 | 0.7826 | 0.0012 | 1e-05 | 128 | 128 | 0.20 | 3 | adam | cosine | none | mean |
| 93 | 0.7692 | 0.0010 | 1e-05 | 128 | 128 | 0.20 | 2 | adam | plateau | none | max |

**Supplementary Table 3 | Full 10-fold cross-validation performance of the top 10 G-LEAP hyperparameter configurations identified by fold 0 screening**

| Rank (10-CV) | Mean AUC ± Std | CV (%) | 95% CI | LR | WD | Batch | Embed | Optimizer | Scheduler | Norm |
|---|---|---|---|---|---|---|---|---|---|---|
| 1 | 0.829 ± 0.028 | 3.38 | [0.8088, 0.8489] | 0.0010 | 1e-05 | 128 | 128 | adam | plateau | batch |
| 2 | 0.821 ± 0.025 | 3.07 | [0.8026, 0.8387] | 0.0008 | 1e-05 | 128 | 128 | adam | cosine | none |
| 3 | 0.820 ± 0.030 | 3.67 | [0.7986, 0.8417] | 0.0008 | 1e-05 | 128 | 256 | adamw | cosine | none |
| 4 | 0.819 ± 0.017 | 2.11 | [0.8069, 0.8316] | 0.0008 | 2e-05 | 128 | 192 | adamw | cosine | none |
| 5 | 0.818 ± 0.028 | 3.43 | [0.7980, 0.8382] | 0.0012 | 1e-05 | 128 | 224 | adamw | cosine | none |
| 6 | 0.817 ± 0.031 | 3.74 | [0.7948, 0.8385] | 0.0010 | 1e-05 | 160 | 128 | adamw | cosine | none |
| 7 | 0.816 ± 0.022 | 2.70 | [0.7998, 0.8313] | 0.0010 | 2e-05 | 128 | 128 | adam | plateau | none |
| 8 | 0.815 ± 0.028 | 3.41 | [0.7954, 0.8352] | 0.0010 | 5e-05 | 128 | 192 | adamw | cosine | none |
| 9 | 0.815 ± 0.022 | 2.71 | [0.7989, 0.8304] | 0.0010 | 5e-05 | 128 | 96 | adamw | cosine | none |
| 10 | 0.814 ± 0.034 | 4.15 | [0.7900, 0.8384] | 0.0012 | 1e-05 | 64 | 128 | adamw | cosine | none |

**Supplementary Table 4 | Performance comparison of G-LEAP and existing methods on GLASS 2.0 dataset under the Random split**

| Method | AUC | Accuracy | Precision | Recall | F1 |
|---|---|---|---|---|---|
| BIND [22] | 0.691±0.038 | 0.680±0.053 | 0.597±0.097 | 0.559±0.112 | 0.574±0.097 |
| DEAttention DTA [23] | 0.948±0.003 | 0.894±0.004 | 0.869±0.007 | 0.859±0.011 | 0.864±0.006 |
| Perceiver CPI [24] | 0.972±0.002 | 0.927±0.003 | 0.905±0.005 | 0.909±0.007 | 0.907±0.004 |
| **G-LEAP (ours)** | **0.975±0.003** | **0.927±0.005** | 0.894±0.007 | **0.921±0.009** | **0.907±0.007** |

**Supplementary Table 5 | Performance comparison of G-LEAP and existing methods on GLASS 2.0 dataset under the Ligand split**

| Method | AUC | Accuracy | Precision | Recall | F1 |
|---|---|---|---|---|---|
| BIND [22] | 0.679±0.037 | 0.686±0.090 | 0.669±0.219 | 0.398±0.277 | 0.440±0.245 |
| DEAttention DTA [23] | 0.940±0.003 | 0.883±0.004 | 0.854±0.013 | 0.850±0.013 | 0.852±0.006 |
| Perceiver CPI [24] | 0.966±0.002 | **0.921±0.003** | **0.898±0.006** | 0.900±0.005 | **0.899±0.004** |
| **G-LEAP (ours)** | **0.969±0.003** | 0.916±0.005 | 0.885±0.006 | **0.902±0.018** | 0.893±0.007 |

### Supplementary figure legends

**Supplementary Figure 1 | Benchmarking of feature encoding strategies and ablation study**

**A,** Performance comparison of stepwise feature refinement across three data splitting strategies (Random, Ligand, and Protein splits) on the initial GLASS dataset. The baseline model uses conventional descriptors (protein mol2vec; compound RDKit descriptors), achieving ROC-AUC scores of 0.706, 0.701, and 0.687 for Random, Ligand, and Protein splits, respectively. Replacing protein features with ESM Cambrian (ESM C) embeddings yielded substantial improvements (Random: 0.797; Ligand: 0.796; Protein: 0.715), demonstrating that evolutionary context captured by protein language models contributes to generalization across both novel compounds and novel receptors primarily through refined protein representations. Further substitution of compound features with Uni-Mol2 embeddings combined with RDKit descriptors resulted in additional gains across all splits (Random: 0.951; Ligand: 0.937; Protein: 0.822). The combination of evolutionary and geometric features consistently yielded the highest ROC-AUC across all splitting strategies, confirming the synergistic benefit of dual-modal feature integration. **B,** Ablation study of the bilinear interaction module. Adding the bilinear interaction module to the GCN backbone consistently improves ROC-AUC compared to simple feature concatenation across all three splitting strategies, demonstrating that the bilinear mechanism effectively captures the non-linear cross-modal dependencies between the protein and ligand latent representations.

**Supplementary Figure 2 | Hyperparameter sensitivity and model stability**

**A,** Hyperparameter heatmaps illustrating the impact of key hyperparameters on model performance (ROC-AUC): embedding dimensions (96–256), learning rates ($8 \times 10^{-4}$-$1.2 \times 10^{-3}$), and weight decay ($5 \times 10^{-6}$–$5 \times 10^{-5}$). The broad high-performance regions (green/yellow zones) indicate that G-LEAP is robust to parameter variations across a wide range of configurations, rather than being dependent on narrowly tuned settings. **B,** Sensitivity analysis of individual hyperparameters. Bar plots show ROC-AUC performance across different batch sizes (left), dropout rates (center), and optimizer choices (right). The orange dashed line indicates the overall mean ROC-AUC (0.814). Performance was comparable across batch sizes from 64 to 160, with a slight decrease observed at 192. Higher dropout rates (0.25–0.30) resulted in reduced performance, and AdamW optimizer showed modestly higher performance than Adam on average. Error bars represent standard deviation across cross-validation folds.

**Supplementary Figure 3 | Family-wise performance of G-LEAP across GPCR classes**

**A–D,** Model performance stratified by GPCR class (Class A, B, C, and F) using held-out test predictions from 10-fold cross-validation. Bar plots report (**A**) Accuracy, (**B**) ROC-AUC, (**C**) Average Precision (AP), and (**D**) F1 score. Class A (Rhodopsin-like) GPCRs, which constitute 96.0% of the dataset (n = 181,192 samples), demonstrated consistently high performance across all metrics (ROC-AUC = 0.801). Performance was lower for data-limited classes: Class B (Secretin; ROC-AUC = 0.693; n = 1,857), Class C (Glutamate; ROC-AUC = 0.655; n = 5,296), and Class F (Frizzled; ROC-AUC = 0.744; n = 275). These variations are expected given the severe class imbalance inherent to current GPCR interaction databases. Notably, G-LEAP maintains meaningful discriminative power (ROC-AUC > 0.65) across all major families, indicating that the model has not overfit to the dominant Class A receptors but retains generalizable recognition capacity across the GPCRome.

**Supplementary Figure 4 | G-LEAP score distributions and ligand retrieval performance for DRD4 and ADORA2A from the DUDE-Z benchmark**

**A**, Distributions of G-LEAP scores for dopamine receptor D4 (DRD4; ligands: n = 37, decoys: n = 1,850) and adenosine A2a receptor (ADORA2A; ligands: n = 90, decoys: n = 4,500) from the DUDE-Z benchmark, comparing known active ligands (orange) and physicochemical decoys (blue). Histograms are normalized to unit area to enable direct comparison between groups despite large differences in sample size. Corresponding results for MTNR1A are shown in Fig.

2E. **B**, Numbers of active ligands retrieved at top-K% thresholds for DRD4 and ADORA2A. Orange bars indicate the observed counts retrieved by G-LEAP, and the green dashed line indicates the number expected under random ranking. The substantial excess of observed over random counts at all top-K thresholds confirms the practical screening utility of G-LEAP for these targets. Corresponding results for MTNR1A are shown in Fig. 2F.

**Supplementary Figure 5 | Large-scale virtual screening of ligand-characterized GPCRs confirms target-specific scoring and chemical space selectivity.**

**A,** Large-scale virtual screening pipeline. The ZINC20 database (9.4 million compounds) was filtered for Lipinski's rule of five [37], PAINS exclusion [38], and Brenk's structural alerts [39], resulting in 6.15 million high-quality drug-like candidates. These compounds were screened against three ligand-characterized GPCR targets: free fatty acid receptor 1 (FFAR1), muscarinic acetylcholine receptor M2 (CHRM2), and orexin receptor type 2 (HCRTR2). **B,** Distribution of G-LEAP prediction scores for FFAR1, CHRM2, and HCRTR2. Histograms display the frequency of prediction scores across the 6.15 million screened compounds for each target. Y-axis values are shown as multiples of $10^6$ (compound count × $10^6$). The distinct, receptor-specific score distributions—with the vast majority of compounds receiving low scores—indicate that G-LEAP infers binding feasibility for only a small, target-specific subset of the chemical library rather than generating spuriously high scores indiscriminately. **C,** Chemical space analysis via UMAP projection. For each GPCR target, compounds were partitioned into four groups: known ligands (red cross marks), top-ranked 1,000 compounds (orange circles), bottom-ranked 1,000 compounds (blue squares), and 1,000 randomly selected compounds (green triangles). UMAP dimensionality reduction based on Morgan fingerprints reveals that high-scoring compounds cluster in chemical space regions more closely resembling known ligands compared to low-scoring and randomly selected compounds.

**Supplementary Figure 6 | G-LEAP retrieves structurally novel ligands demonstrating scaffold-hopping capacity.**

**A**, Reference similarity distributions for novelty threshold definition. Histograms show pairwise Tanimoto similarity scores computed using three molecular fingerprints (Morgan FP, RDKit FP, and MACCS keys) for 499,500 unique pairs generated from 1,000 randomly retrieved compounds from the PubChem database (accessed January 7, 2026). For each fingerprint, the mean similarity and the 99th percentile (top 1%) are indicated. These distributions were used to define fingerprint-specific novelty cutoffs: compounds with Tanimoto similarity below the indicated thresholds are considered structurally distinct from the reference set. **B**, Distribution of structural novelty for top-ranked G-LEAP candidates. For each GPCR target—FFAR1, CHRM2, and HCRTR2—histograms show the distribution of maximum Tanimoto similarity to all known active ligands for each top-1,000 candidate compound, computed with Morgan FP (left), RDKit FP (middle), and MACCS keys (right). Red dashed lines denote the novelty cutoffs established in panel A (Morgan: 0.25, RDKit: 0.50, MACCS: 0.65). The x-axis indicates maximum similarity to known actives and the y-axis indicates compound count. A substantial proportion of high-scoring compounds fall below these cutoffs across all three fingerprint types and all three targets, demonstrating that G-LEAP successfully identifies structurally novel scaffolds markedly distinct from established ligands, confirming robust scaffold-hopping capacity.

**Supplementary Figure 7 | Robustness of metabolome-scale GPCR screening and validation of prediction mechanisms**

**A,** Fold-wise ROC curves for the ensemble model used in HMDB screening. Performance consistency across the 10-fold cross-validation models demonstrates stable generalization to unseen GPCRs. The tight clustering of individual fold curves (colored lines) around the mean ROC curve (bold line) indicates that the ensemble approach mitigates overfitting to any single data partition, ensuring robust predictions during atlas construction. **B,** Target fishing validation ("Reverse Screening"). Rank distribution of known target receptors when promiscuous endogenous metabolites (serotonin, dopamine, acetylcholine) are screened against the entire GPCR superfamily (n = 624 receptors). Box plots show the percentile rank of known targets; 78% of known GPCR partners ranked within the top 10% (mean percentile: 7.1%), demonstrating G-LEAP's utility for identifying polypharmacological interactions in a bidirectional manner (ligand → target). **C,** Chemical diversity of top-100 predictions. Distribution of maximum Tanimoto similarity coefficients between top-100 HMDB metabolites predicted by G-LEAP and known endogenous ligands for each GPCR. Green and orange lines denote the median (0.086) and mean (0.111), respectively. The low global similarity (median = 0.086) confirms that G-LEAP identifies chemically diverse candidate ligands beyond simple nearest-neighbor retrieval based on structural resemblance, indicating that the model captures deeper compatibility signals such as protein-sequence patterns and learned interaction motifs. **D,** Receptor-specific structural similarity of false positives. For each GPCR, bar length indicates the mean Tanimoto similarity between top-100 non-annotated predictions (false positives; compounds not recorded as known ligands in GtoPdb) and the true known endogenous ligand. Percentages displayed to the right of each bar indicate the fraction of these 100 false positives exhibiting moderate-to-high structural similarity (Tanimoto > 0.3) to the true ligand. For sphingosine-1-phosphate receptors (S1P1, S1P3, S1P5) and adenosine A3 receptor, 40–63% of high-scoring non-annotated compounds exhibit moderate structural similarity (Tanimoto > 0.3) to the known endogenous ligand, indicating that the model preferentially ranks compounds sharing substructural features with true ligands for these receptors.

**Supplementary Figure 8 | Tissue co-expression validation of predicted orphan GPCR-ligand pairs**

**A, B**, Tissue-specific co-expression analysis using GTEx v10 RNA-seq data across 54 human tissues. Scatter plots show the correlation between orphan receptor expression and the expression of biosynthetic enzymes responsible for producing the predicted endogenous ligand. (**A**) *GPR20* versus sphingosine kinase 1 (*SPHK1*), the enzyme that phosphorylates sphingosine to generate sphingosine 1-phosphate [54]. Spearman correlation: $\rho = 0.627$, $p < 10^{-8}$. **(B)** *GPR26* versus arylalkylamine *N*-acetyltransferase (*AANAT*), the rate-limiting enzyme in melatonin biosynthesis [55]. Spearman correlation: $\rho = 0.339$, $p < 0.01$. These significant positive correlations reinforce the physiological plausibility of the predicted orphan receptor–ligand pairs, as co-expression with biosynthetic machinery provides independent transcriptomic evidence supporting functional coupling in human tissues.

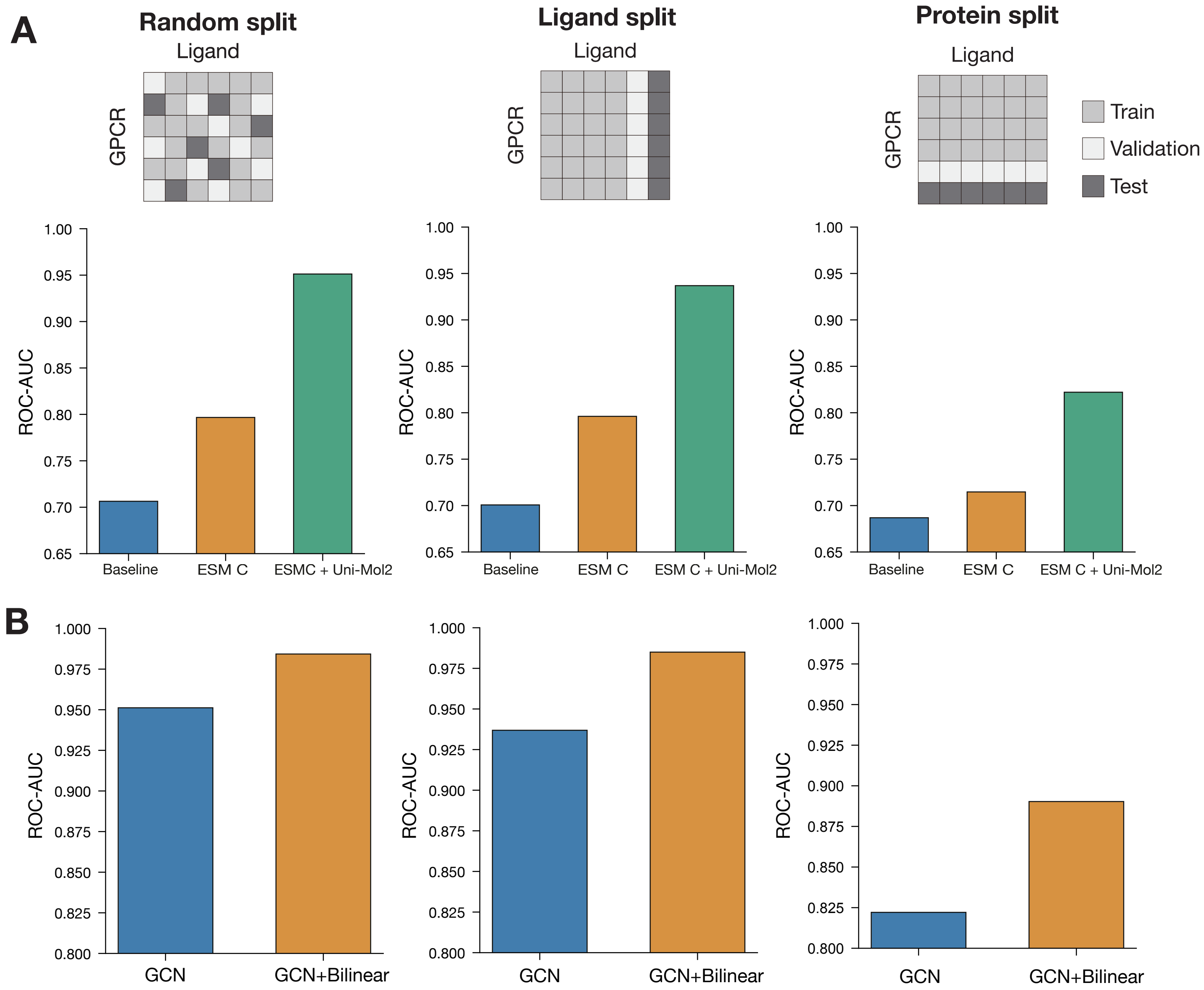


Sakuma et al., Figure S1

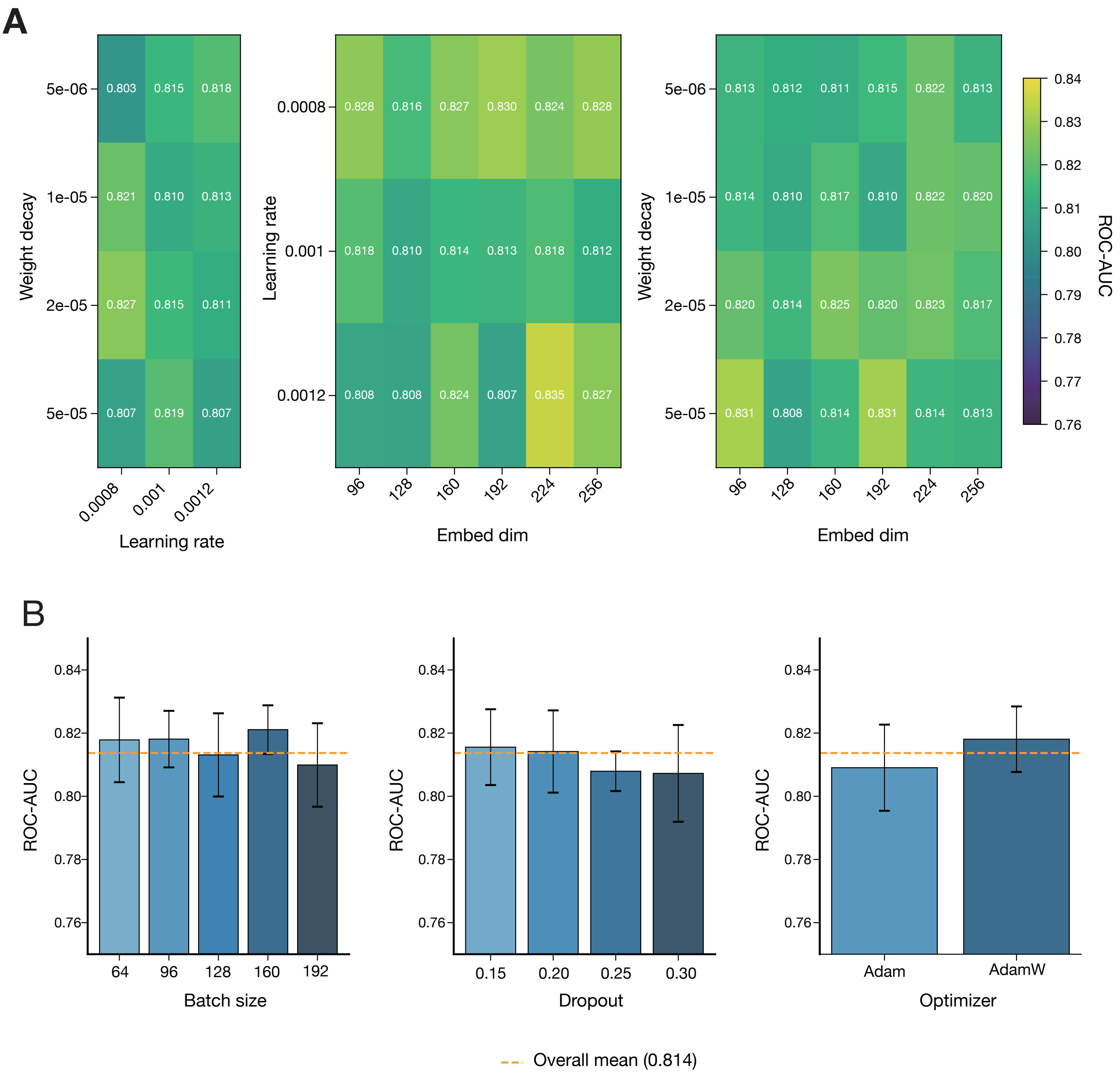


**Sakuma et al., Figure S2**

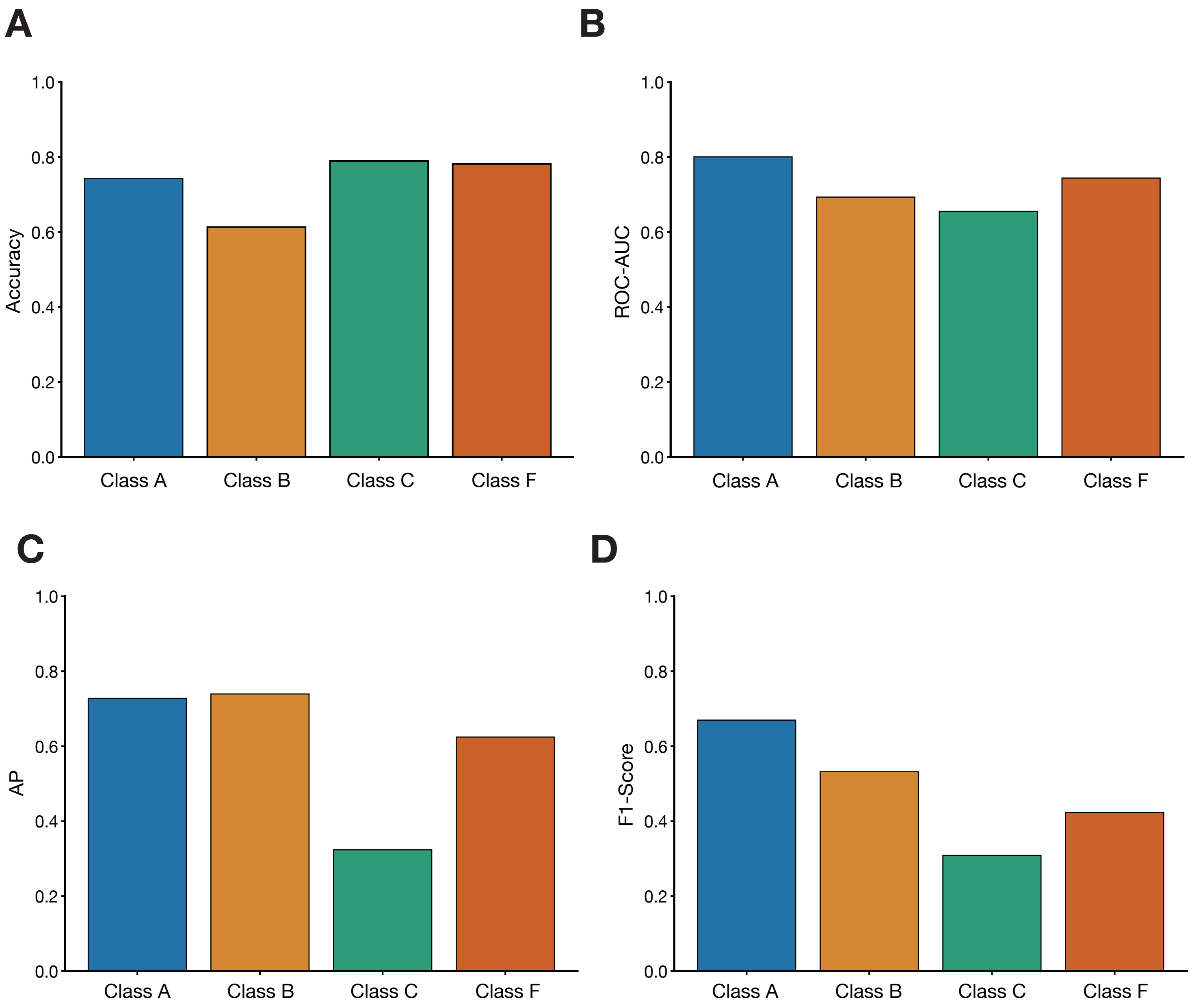


**Sakuma et al., Figure S3**

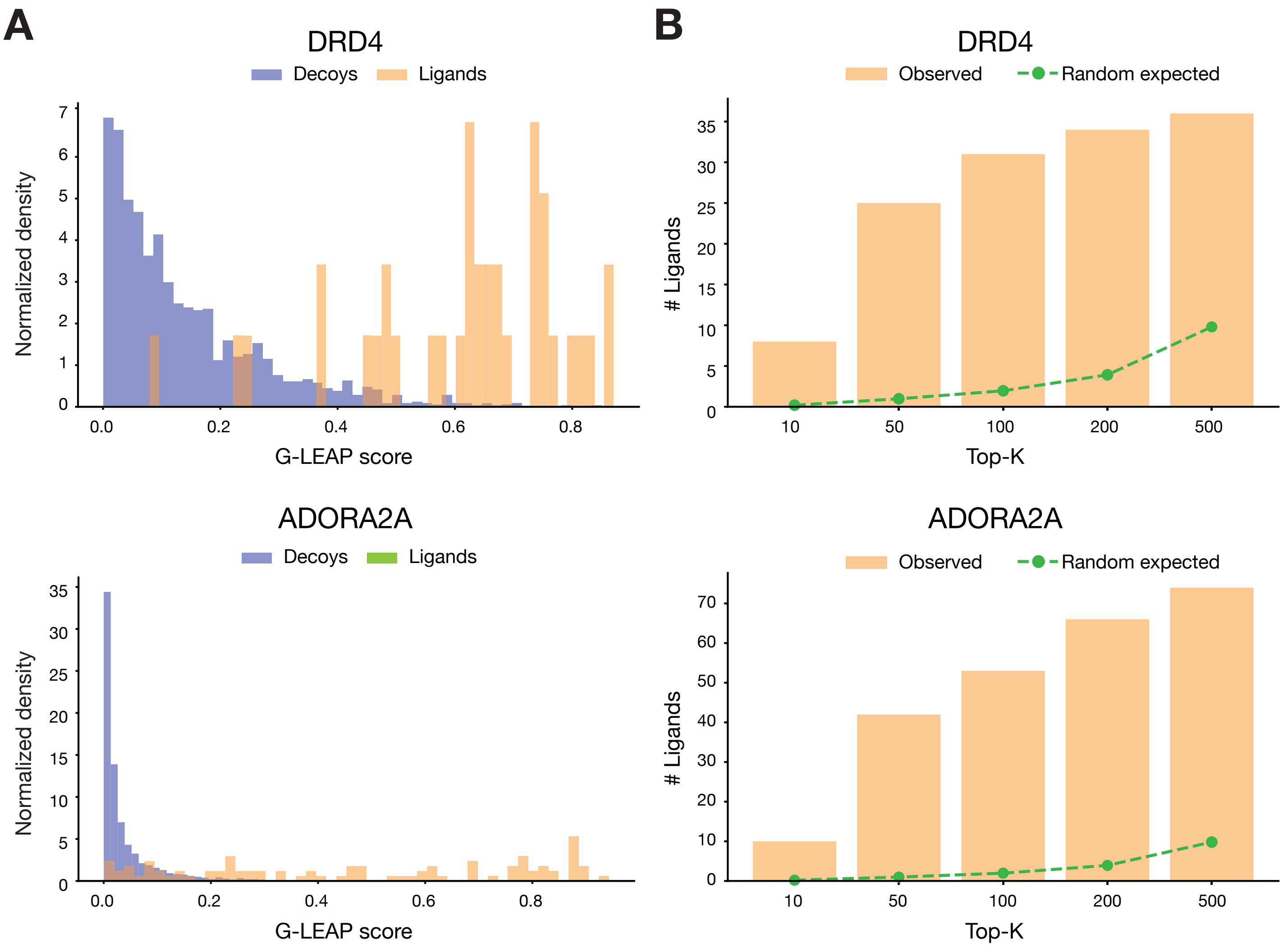


**Sakuma et al., Figure S4**

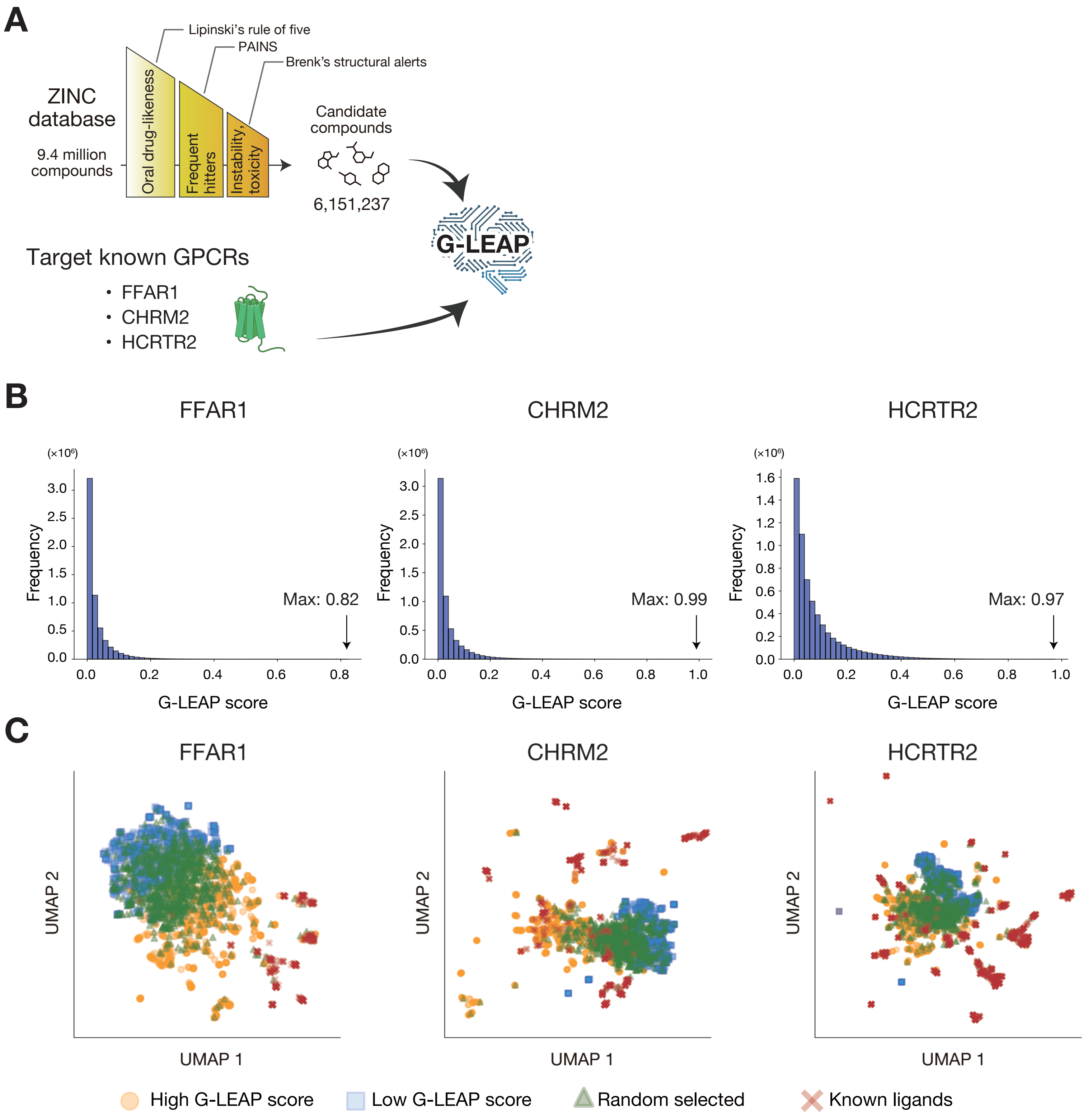


**Sakuma et al., Figure S5**

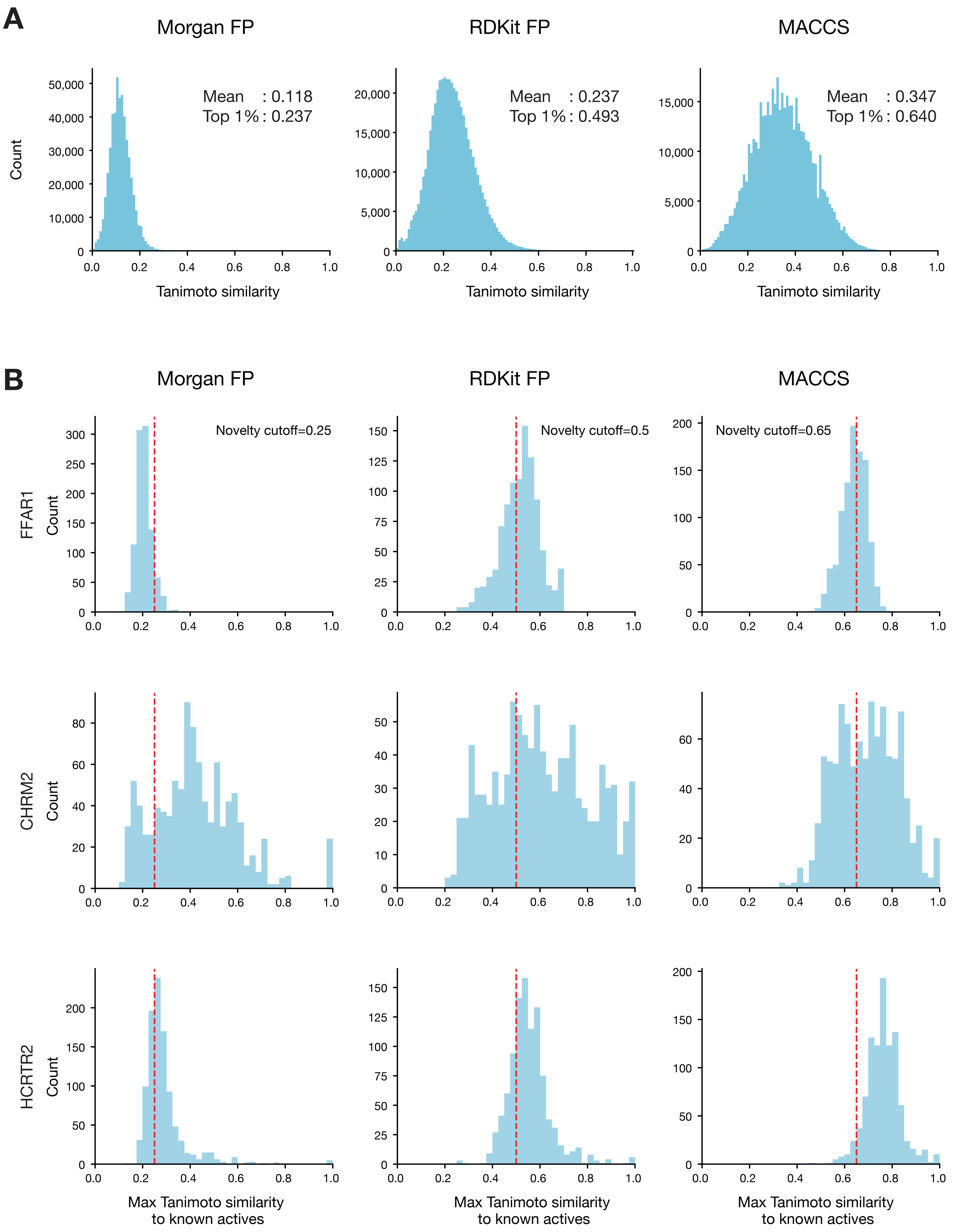


**Sakuma et al., Figure S6**

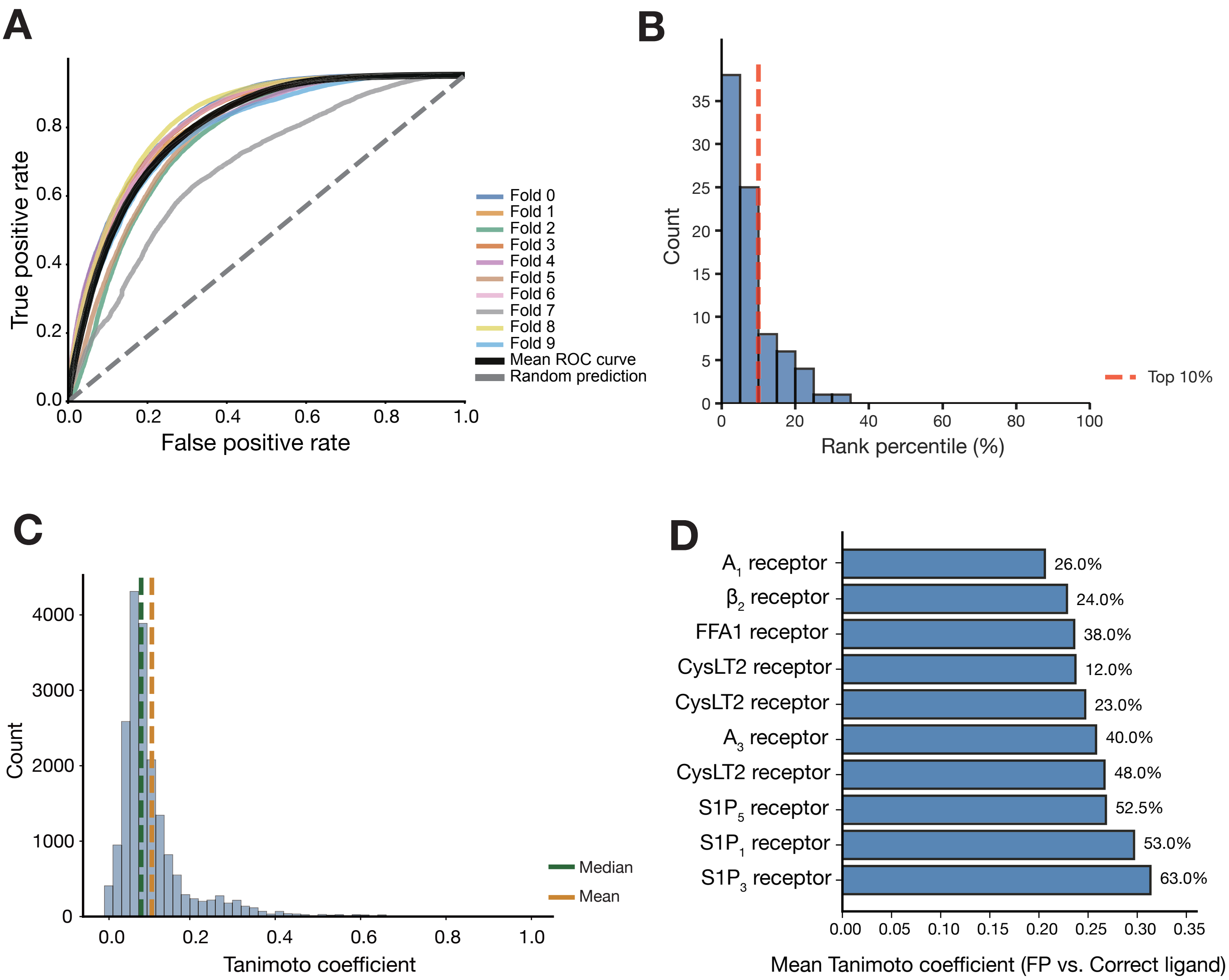


**Sakuma et al., Figure S7**

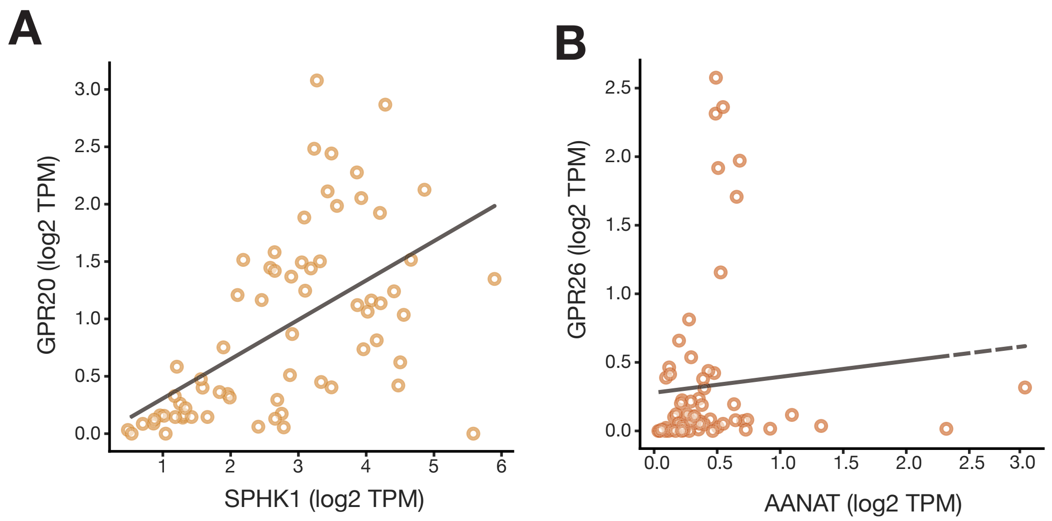


**Sakuma et al., Figure S8**
